# A Single Cell Time Course of Senescence Uncovers Discrete Cell Trajectories and Transcriptional Heterogeneity

**DOI:** 10.1101/2023.02.17.529001

**Authors:** Serban Ciotlos, Lauren Wimer, Judy Campisi, Simon Melov

## Abstract

Senescent cells (SnCs) are typically studied as endpoints of a complex transformational process, owing to their frequent maladaptive effects on surrounding tissue and cells. SnCs accumulate with age, and while they ultimately comprise a small percentage of cells in tissues, they have important roles in age associated pathologies. Several obstacles remain in understanding the heterogeneous nature of senescence, and formulating potent beneficial intervention strategies. One approach targets senescent cells and kills them (“senolytic” approach), and is often driven by a low resolution understanding of SnC identity, which risks both incomplete clearance and off-target effects. Cellular senescence is not a singular binary response, but a suite of response trajectories that vary by multiple parameters including inducer and initial cell state. In order to elucidate the developmental trajectories of SnCs, we performed single-cell RNA sequencing on IMR90 lung fibroblasts senescencing across a 12 day time period. Our analysis reveals substantial heterogeneity in gene expression within timepoints and across the full time-course. We uncovered unique markers and differentially regulated pathways in cell populations within each timepoint. Supervised trajectory inference of the time-course data uncovered the root-origin and fates of distinct SnC lineages over 3 stages of senescence induction. Altogether our data provide a novel approach to stud SnC development, identifying cell states of interest, and differentiating between SnCs and quiescent cells. This will aid in identifying key targets for therapeutic intervention in senescence.

## Introduction

Cellular senescence is a complex cell state characterized by permanent cell cycle arrest, deregulated metabolism, macromolecular damage, and an altered secretory profile relative to cycling or quiescent cells. Multiple types of stimuli can provoke cellular senescence and a cell type-specific senescence-associated secretory phenotype (SASP) [1–4]. SnCs are implicated in many physiological processes (both adaptive and maladaptive) and diseases of aging, including Alzheimer’s and Parkinson’s disease, atherosclerosis and cardiovascular dysfunction, chemotherapy cardiotoxicity, and osteoarthritis. Benign processes associated with senescence include wound healing, tissue regeneration, embryogenesis, and tumor suppression [5–18]. Senescence is triggered by multiple distinct inducers including telomeric and non-telomeric DNA damage (from genotoxic drugs such as the chemotherapy drug doxorubicin), irradiation, oncogenic stress, oxidative stress, and mitochondrial dysfunction [19]. The inducer chosen for specific cell types is a source of potential heterogeneity in the senescence response. In this study we induced human fetal lung fibroblasts (IMR90) to senesce using doxorubicin (doxo), a chemotherapy drug that has multiple deleterious side effects including cardiotoxicity and induction of senescence. Two major pathways have been heavily investigated in the context of senescence: the p53/p21 (p21/CDKN1A) and p16Ink4a/retinoblastoma (p16/CDKN2A) protein pathways. Across species and cell types, there is variability in the senescence response [20]. Many inducers activate either or both p21 and p16 [21]. SnCs acquire a SASP, which consists of chemokines, cytokines, growth factors, and matrix remodeling proteins [22–24]. The SASP is a pro-inflammatory profile that leads to alterations in tissue microenvironment and paracrine transmission of cell senescence [2, 25–27]. Major components of several SASPs include interleukins IL-1, 6, and 8, vascular endothelial growth factor A (VEGFA), and matrix metalloproteinases such as MMP3. Proinflammatory cytokine components of the SASP, such as IL6, IL8, IL1α/β, TNFα, and TNFSF7 are generally conserved across senescence contexts. However, as with senescence markers in general, the composition of the SASP is fluid, and varies by induction method, cell type, and organism [22, 24, 28–31]. Heterogeneity in cellular senescence refers to the variation in the molecular and functional responses of senescent cells, despite their shared phenotype of growth arrest [20]. This variability contributes to the diverse roles that SnCs can play in physiological processes and age-related disorders. The killing or modification of SnCs has been shown to ameliorate age related pathologies and extend lifespan in mice [6]. Several approaches can be used to accomplish this, including senolytic molecules, immune modulation, and “senomorphic” compounds which suppress the detrimental aspects of the SASP without killing the cell [32]. Current therapeutic strategies are based on a limited understanding of senescence, often targeting non-specific molecules or pathways that are shared broadly across the SnCs of different cell types. For example, resistance to apoptosis is a shared feature of SnCs. The drug Navitoclax (ABT-263) inhibits anti-apoptotic protein BCL2 and induces SnCs to undergo apoptosis [33]. However, this method also risks eliminating beneficial or neutral cells. Transgenic mouse models of SnC clearance such as the p16-3MR line also depend on a broad targeting strategy, whereby Ganciclovir kills any cell expressing a p16-driven reporter construct. This approach risks eliminating non-senescent cells [12]. For example, macrophages express p16 as a normal response to immune stimuli, not senescence induction [34]. In addition, this paradigm will not target SnCs lacking p16 expression, as the method relies on defining SCs by p16 expression alone. Previous work on SnC heterogeneity (at the bulk RNA level) showed that the transcriptional response to induction changes across time, while noting that the ability to discriminate between senescence subtypes is critical to developing specific therapy targets [20, 35]. In order to identify specific and time-sensitive gene markers of SnCs, we used single cell RNA sequencing to survey and compare untreated proliferating, quiescent, and senescing cells across 10 timepoints derived from a single culture stock. A population of senescence-evading cells was identified across the time course, which resisted both initial senescence induction, and remained proliferative despite being cultured in a SASP-enriched environment. These cells distinguished themselves transcriptionally from both normal proliferating cells and stable SnCs. We observed that p16 and p21 expression was not limited to SnCs, and also identified specific markers for each of the 3 main cell states that were studied. Finally, we performed pseudotime and trajectory analyses, uncovering transcriptionally distinct lineages of SnCs and populations engaged in different senescence associated damage/stress responses, cytokine signaling patterns, and metabolic activities.

## Results

To explore the heterogeneity of cellular senescence both as an end state and a dynamic process, we cultured embryonic lung fibroblasts (IMR-90) in three cell states: proliferation, quiescence, and senescence. To induce quiescence, we cultured cells in serum-starved media; for senescence induction we treated cells with doxo for 24h and harvested them at 12 hours, 1 day, 2, 4, 6, 8, 10, or 12 days post-dose (Figure 1A) for single cell profiling. These cell states and time points were chosen to capture transcriptional changes as cells transform from a proliferative state to a senescent state. To ensure robustness in our experimental design, all experiments were run with two biological replicates. Single cells were prepared by microfluidic encapsulation and libraries were prepared according to the 10x protocol and submitted for high throughput sequencing at the UC Davis Core Sequencing facility. Quality control included the removal of cells with low gene and read counts, those that were highly enriched for mitochondrial and ribosomal gene expression, as well as those which exhibited long non-coding RNAs MALAT1 and NEAT1. QC exclusion resulted in 53,435 cells from Replicate 1 and 79,074 cells from Replicate 2 (Figure 1B) with an average sequencing depth of 13,251 reads per cell and 11,726 reads per cell, respectively. Average gene counts were 2,972 and 2,966 for Replicate 1 and Replicate 2, respectively. Replicates were visualized together to assess whether significant differences existed in cell embedding and clustering (Figure1C), which could indicate either biological or technical variation. Although the replicates do not overlap perfectly, we did not observe strong batch effects necessitating correction, a process which may reduce biological variation while attempting to correct for technical variation. Cells from each timepoint do not strongly segregate by replicate, indicating batch effects did not significantly affect library quality. We determined that the percent of reads assigned to cells declined at D6, D8, and D10 post-doxo before slightly recovering at D12 (Figure 1C). This decrease is due to an increase in ambient (non-cell-assigned) RNA, suggesting that a fraction of senescing cells become fragile 6-10 days following doxo treatment, and may prematurely lyse due to the shear stress that cells experience during microfluidic processing. Therefore, these cells were excluded from our analysis. Due to the high degree of homogeneity between replicates, we combined them for increased power to characterize heterogeneity in a sizable cell population.

**Figure 1.**
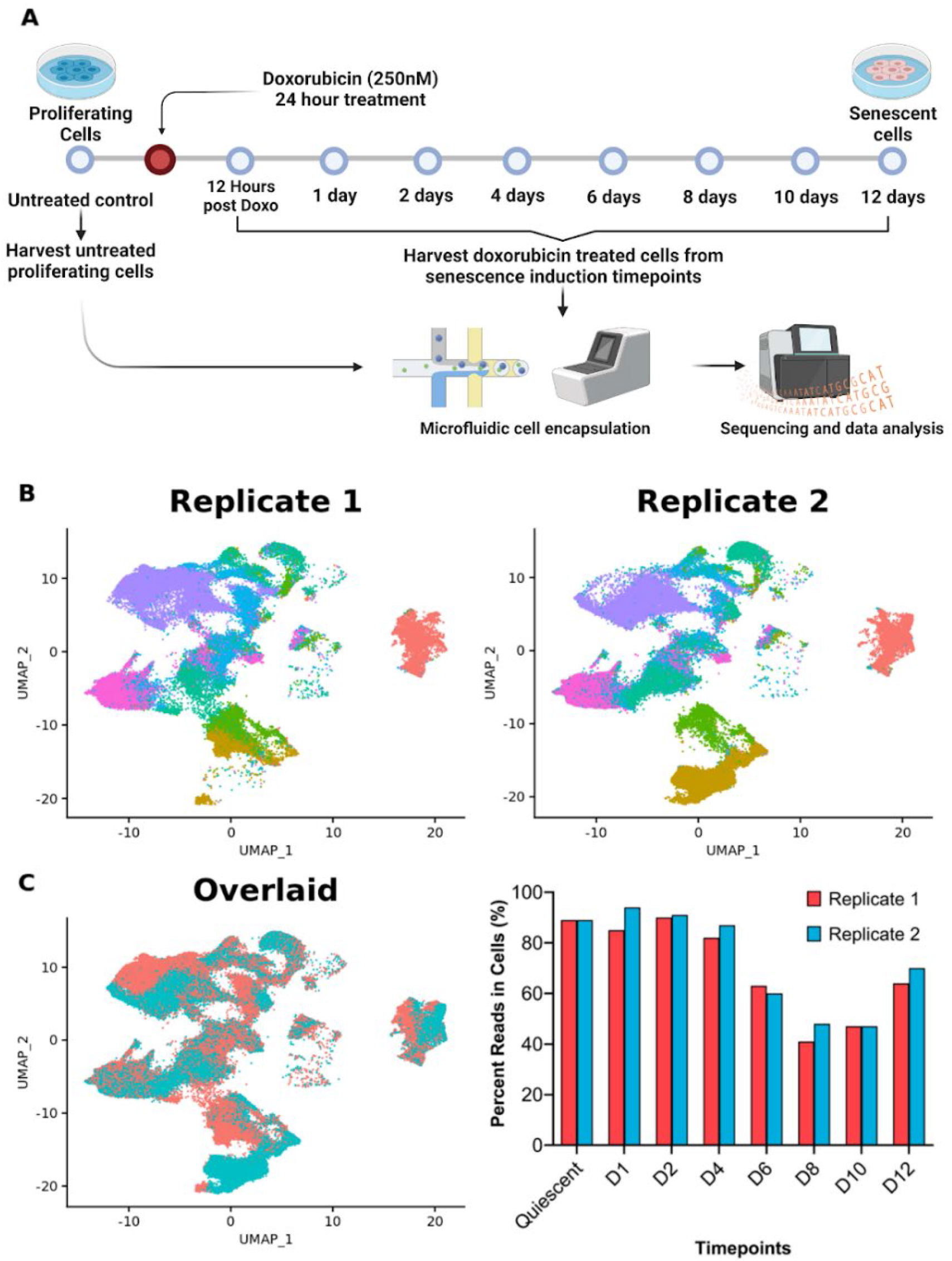
Experimental overview for inducing cellular senescence and quality control of replicate heterogeneity. (A) Proliferating IMR90 cells treated with 250 nM doxo for 24h are shown at either 12 hours (referred to as “H12”),1 (“D1”), 2 (“D2”), 4 (“D4”), 6 (“D6”), 8 (“D8”), 10 (“D10”), or 12 (“D12”) days post induction. (B) Timepoint replicates visualized by UMAP. (C) Percent of reads assigned to validated cells from each timepoint replicate library.

### Proliferating cells co-cultured with senescing cells

Despite treatment with doxo, we observed that some cells were able to “evade” senescence conversion. Including untreated proliferating cells, we identified 8,449 cells across the 12-day time course which remained proliferative (expressing MKI67+/CCNA2+/TOP2A+/MCM2+/PCNA+); we visualized these cells at each time point using UMAP (Figure 2). The presence of proliferating cells suggests that not all cells within a culture are induced to senescence following 24h of doxo treatment. To further characterize these cells, we identified differentially expressed genes and upstream regulators, between sequential timepoints (Table 1). 12 hours after doxo treatment of proliferating cells, the growth factor binding protein IGFBP5 and matrix metallopeptidase MMP1 are the top up- and down-regulated genes. We identified ILF3, a transcription factor known to also bind mRNAs of SASP proteins, and TP53, a central activator of the senescence response, as activated upstream regulators. Conversely, the activity of the transcription factor E2F3 and methyltransferase DNMT3B was inhibited. From 12-24 hours (H12-D1), we observed increases in growth factor and the SASP protein GDF15, and decreases in another SASP factor, SFRP1. TP53 gene expression activity continues to increase from H12 to D1, and histone demethylase KDM5B gene expression changes, consistent with its epigenetic function. Transcription factors FOXM1, which plays important roles in aging and senescence, and HSF1 (heat shock regulator whose suppression activates p38-NF-κB-SASP) are inhibited. Comparing those cells that escape senescent transformation from D2 to D1, we determined that IGFBP5 is upregulated. In contrast to the previous (D1 vs. H12) comparison, GDF15 is strongly downregulated by D2. The reduction in GDF15 may indicate an immediate response that is attenuated with time, or that such cells are able to suppress the need to release GDF15. Gene expression of JARID2 and BACH1 is also increased. The latter is a known inhibitor of p53 and oxidative stress-induced senescence. Indeed, transcriptomic regulation by TP53 is inhibited at D2, reversing the trend of the previous time points. Histone deacetylase HDAC3 function, recently shown to repress p65 NF-κB-mediated induction of the SASP, is also inhibited. Between D2 and D4, expression of the known SASP factor CCL2 is increased, while gene expression of the secreted factor and chaperone CLU is downregulated. This finding illustrates the unique transcriptional activity that separates the senescence resistant cells from more robust SnCs at this time point.

**Figure 2:**
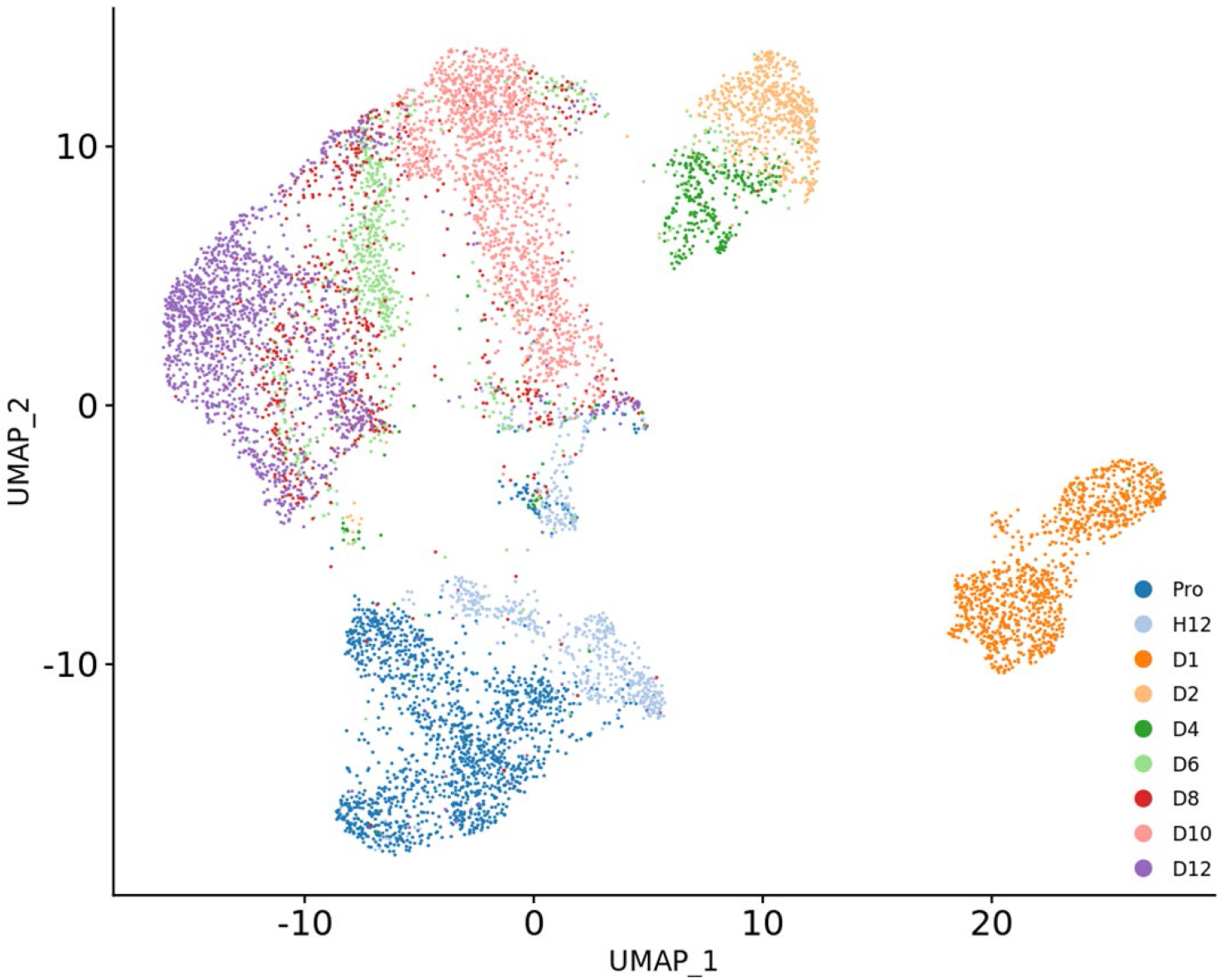
Visualization of proliferating cells. UMAP of 8,449 proliferating cells, drawn from both the untreated proliferating (“Pro”) timepoint and senescence resistant (H12-D12) timepoints, identified across the full time course.

**Table 1.**
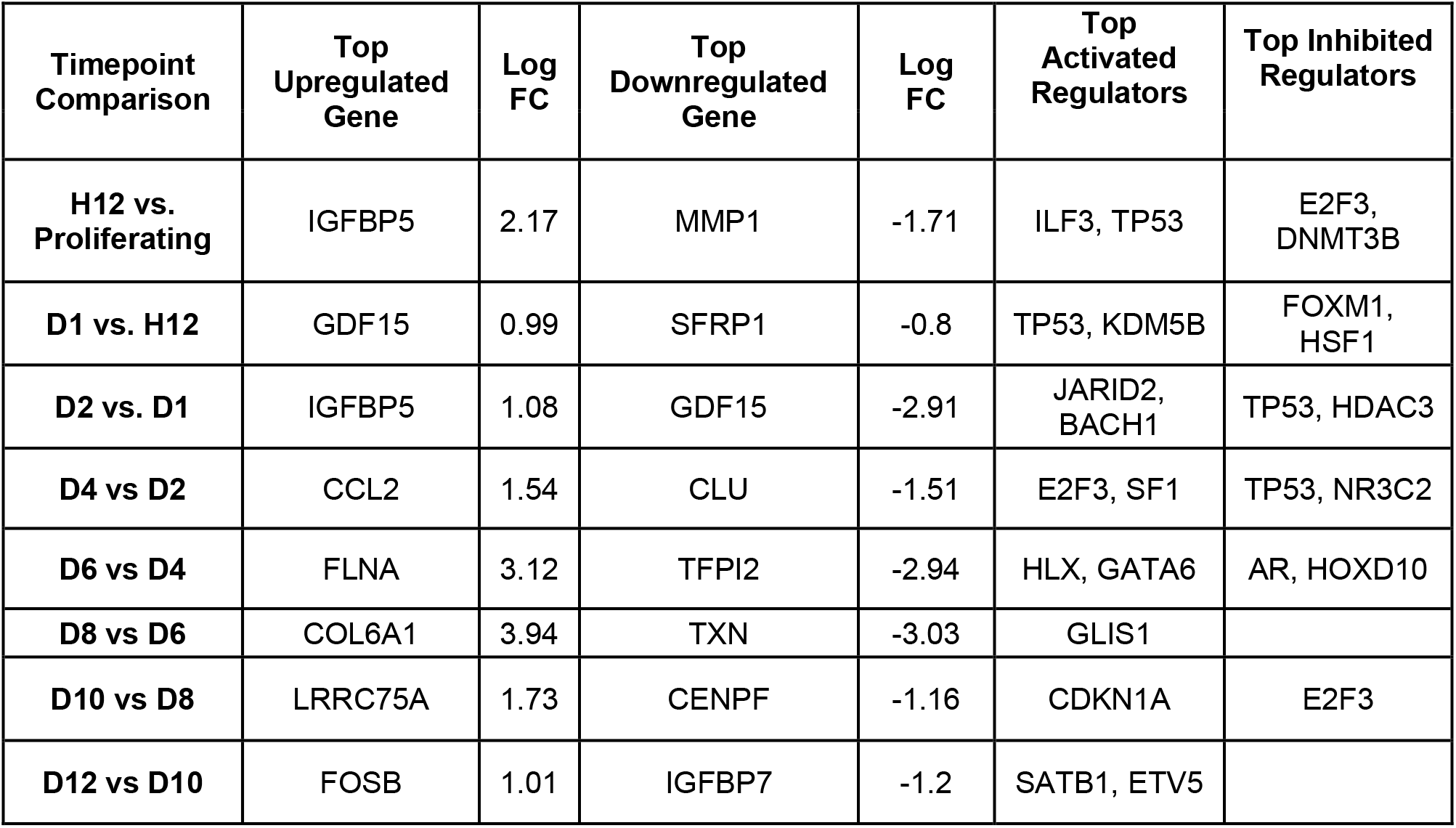
Top differentially expressed genes and transcriptional regulators between senescence resistant cells from sequential timepoints. Differentially expressed genes (DEGs) were identified with Seurat’s FindMarkers function, and regulators were identified with Ingenuity IPA. Empty boxes indicate a result flagged for bias by Ingenuity’s IPA analysis, and are not reported.

| Timepoint Comparison | Top Upregulated Gene | Log FC | Top Downregulated Gene | Log FC | Top Activated Regulators | Top Inhibited Regulators |
| --- | --- | --- | --- | --- | --- | --- |
| **H12 vs. Proliferating** | IGFBP5 | 2.17 | MMP1 | -1.71 | ILF3, TP53 | E2F3, DNMT3B |
| **D1 vs. H12** | GDF15 | 0.99 | SFRP1 | -0.8 | TP53, KDM5B | FOXM1, HSF1 |
| **D2 vs. D1** | IGFBP5 | 1.08 | GDF15 | -2.91 | JARID2, BACH1 | TP53, HDAC3 |
| **D4 vs D2** | CCL2 | 1.54 | CLU | -1.51 | E2F3, SF1 | TP53, NR3C2 |
| **D6 vs D4** | FLNA | 3.12 | TFPI2 | -2.94 | HLX, GATA6 | AR, HOXD10 |
| **D8 vs D6** | COL6A1 | 3.94 | TXN | -3.03 | GLIS1 |  |
| **D10 vs D8** | LRRC75A | 1.73 | CENPF | -1.16 | CDKN1A | E2F3 |
| **D12 vs D10** | FOSB | 1.01 | IGFBP7 | -1.2 | SATB1, ETV5 |  |

Changes in gene expression between the next pair of timepoints (from D4 to D6) describe a transcriptomic profile more akin to senescent cells: actin-binding protein FLNA is upregulated and serine protease inhibitor TFPI2 is downregulated. TFPI2 is an inhibitor highly expressed in non-invasive cells, and enriching for TFPI2 increases apoptosis and senescence. FLNA was shown to induce senescence in the context of hypoxia, by complexing with DRP1 and leading to mitochondrial fragmentation. At this point, gene expression of the senescence-induced GATA6 transcription factor is increased, while androgen receptor (AR) and HOXD10 activity is diminished. COL6A1 is the top upregulated gene in senescence “evaders” (SnEs) from D6 to D8, confirming previous data indicating that its knockdown suppress cell proliferation and migration. TXN gene expression is downregulated; its suppression has previously been shown to induce senescence in normal human fibroblasts. Expression of the transcription factor GLIS1 is activated, suggesting a link between senescence resistance and its roles as a facilitator of pluripotency induction in somatic reprogramming and fibrosis of senescent fibroblasts. Between D8 and D10, LRRC75A and CENPF are respectively up- and downregulated. The former is a long noncoding RNA whose associated transcripts play multiple roles in regulating proliferation and senescence. Interestingly, its overexpression in spontaneous pre-term birth is associated with an increase in p53/p21 signaling and secretion of SASP factors IL-8 and TNF-α. CENPF is required for chromosomal segregation and kinetochore function. CDKN1A/p21 activity is increased at this time point, potentially indicating a different role for this gene in senescence-resistant cells and SnCs. These cells are ‘proliferative’ but resemble transcriptomic vignettes of senescent cells. FOSB is the top upregulated gene from D10 to D12, consistent with previous findings linking its expression to increased proliferation. IGFBP7 is secreted by SnCs and its upregulation inhibits proliferation; in our study this gene is downregulated. Transcription factors SATB1 and ETV5 are both increased in gene expression at this time point, which supports previous findings indicating their loss suppresses proliferation. Overall these findings describe a distinct and dynamic transcriptional profile for cells that evade doxo-induced senescence while also resisting the impact of SASP-conditioned media. Alongside this longitudinal progression, we found that senescence evaders consistently modulate (either by activation or inhibition) some pathways at every single time point, when compared to their growth arrested neighbors (Table 2). Most pathways involved in cell cycle regulation and chromosomal replication are activated, however specific exceptions (ex. G2/M DNA damage checkpoint regulation) are inhibited. Perhaps most strikingly, both Nucleotide excision and Base excision repair mechanisms are consistently activated, indicating the proliferative capacity of senescence evaders involves a sustained DNA repair response. Several immune-related pathways are involved as well, such as activation of phagocytosis mediated by Fcγ and the immunogenic cell death signaling pathway. In contrast, granzyme A signaling is repressed, indicating that senescence evaders are not priming for caspase-independent apoptosis. Lastly, HMGB1 and VEGF signaling are upregulated, both processes consistent with a proliferative state. These findings together suggest that senescence evading cells walk a fine line between proliferation, apoptosis, and senescence.

**Table 2.**
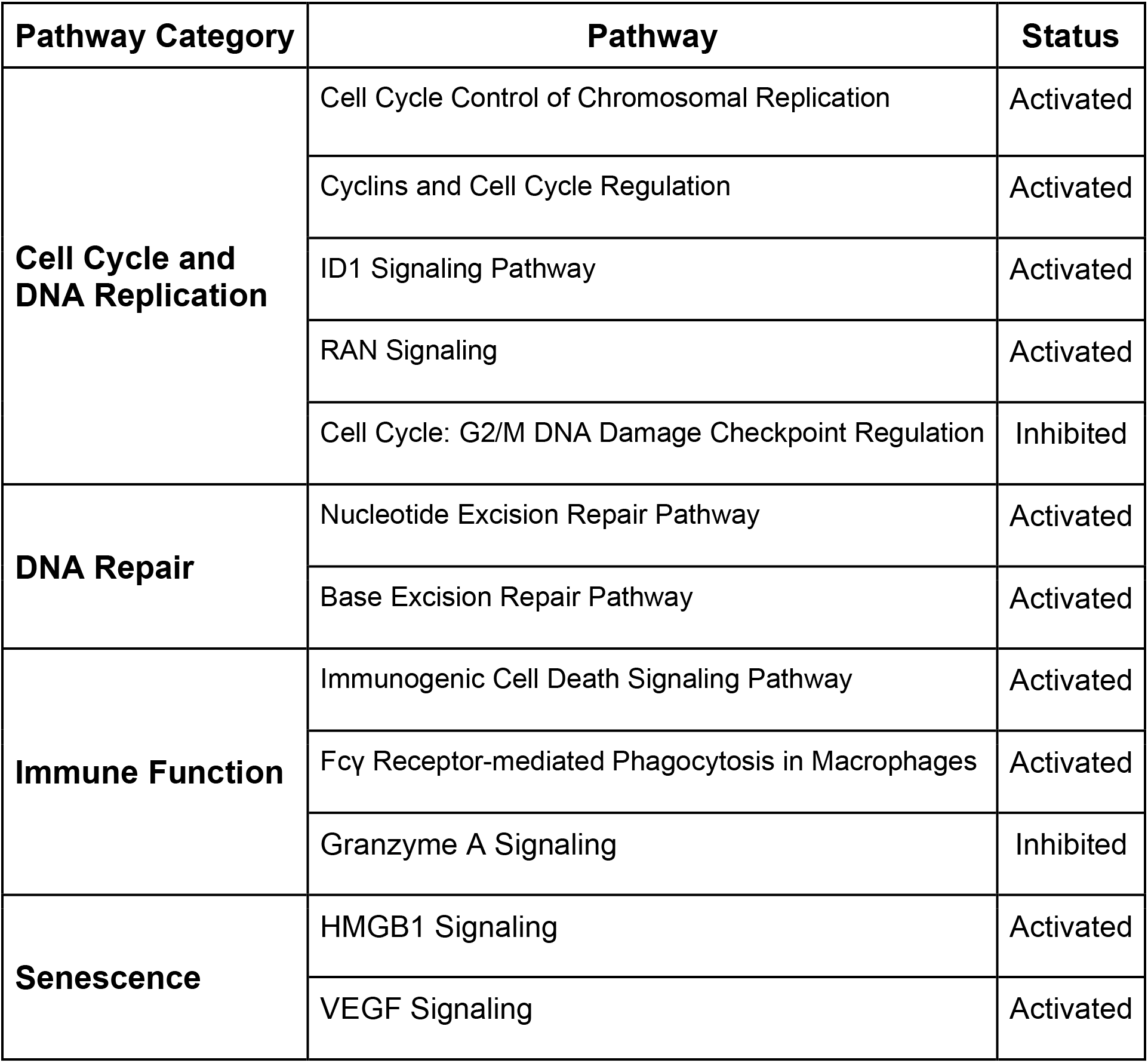
Pathway analysis of activated and inhibited pathways in senescence-resistant cells compared to senescent cells. Consistently modulated pathways of both proliferating and senescing cells at all timepoints. “Activated” denotes higher pathway activity in proliferating cells while “Inhibited” denotes lower pathway activity in proliferating cells. Analysis was performed by Ingenuity’s IPA analysis.

### Differentiating between proliferating, senescent, and quiescent cells

We compared the 3 main outcomes from doxo treatment (proliferating, quiescent and senescent cells; 31,444 cells total) in order to define the transcriptional differences between cell states. Here we report transcriptional markers for each cell state, while also highlighting heterogeneity within respective cell states that may otherwise confound identification. Some cells from the D12 time point retain proliferative capacity (Figure 3A, top), as previously discussed. These cells fall into cluster 12 (Figure 3A, bottom), and are not included in the results here. A small number of quiescent cells clustered (cluster 14) in close spatial proximity to the main body of untreated proliferating cells (clusters 10 and 11). In the (untreated) proliferating cell time point, MKI67 was not ubiquitously expressed (found in 45% of cells), nor were other proliferation markers such as TOP2A (43%), CCNA2 (36%), and MCM2 (25%) (Figure 3B). Proliferating cell nuclear antigen (PCNA) was expressed in 62% of these cells, as opposed to 55% of QnCs and 16% of SnCs. In the (untreated) proliferating cell time point, MKI67 was not ubiquitously expressed (found in 45% of cells), nor were other proliferation markers such as TOP2A (43%), CCNA2 (36%), and MCM2 (25%) (Figure 3B). Proliferating cell nuclear antigen (PCNA) was expressed in 62% of these cells, as opposed to 55% of QnCs and 16% of SnCs.

**Figure 3:**
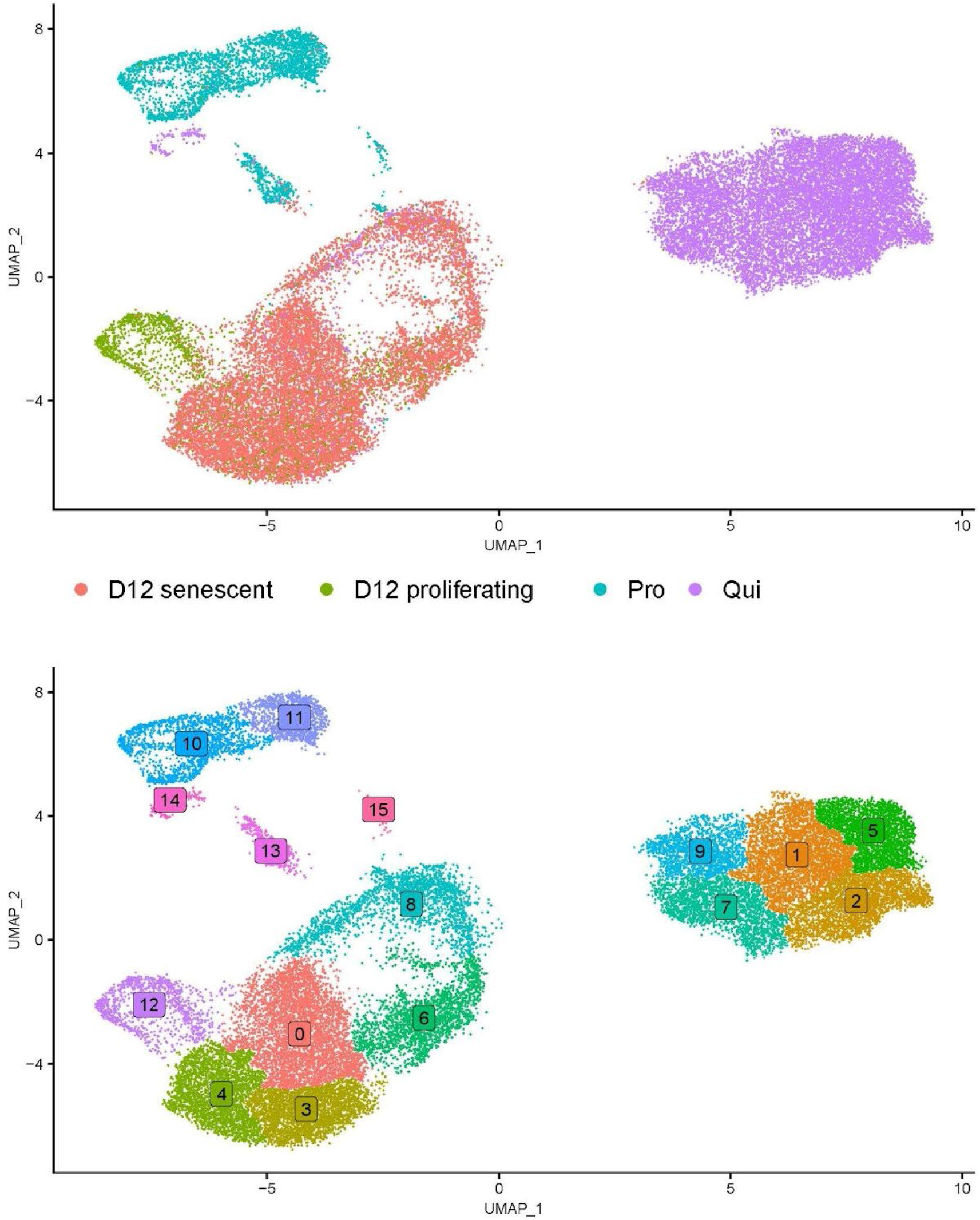
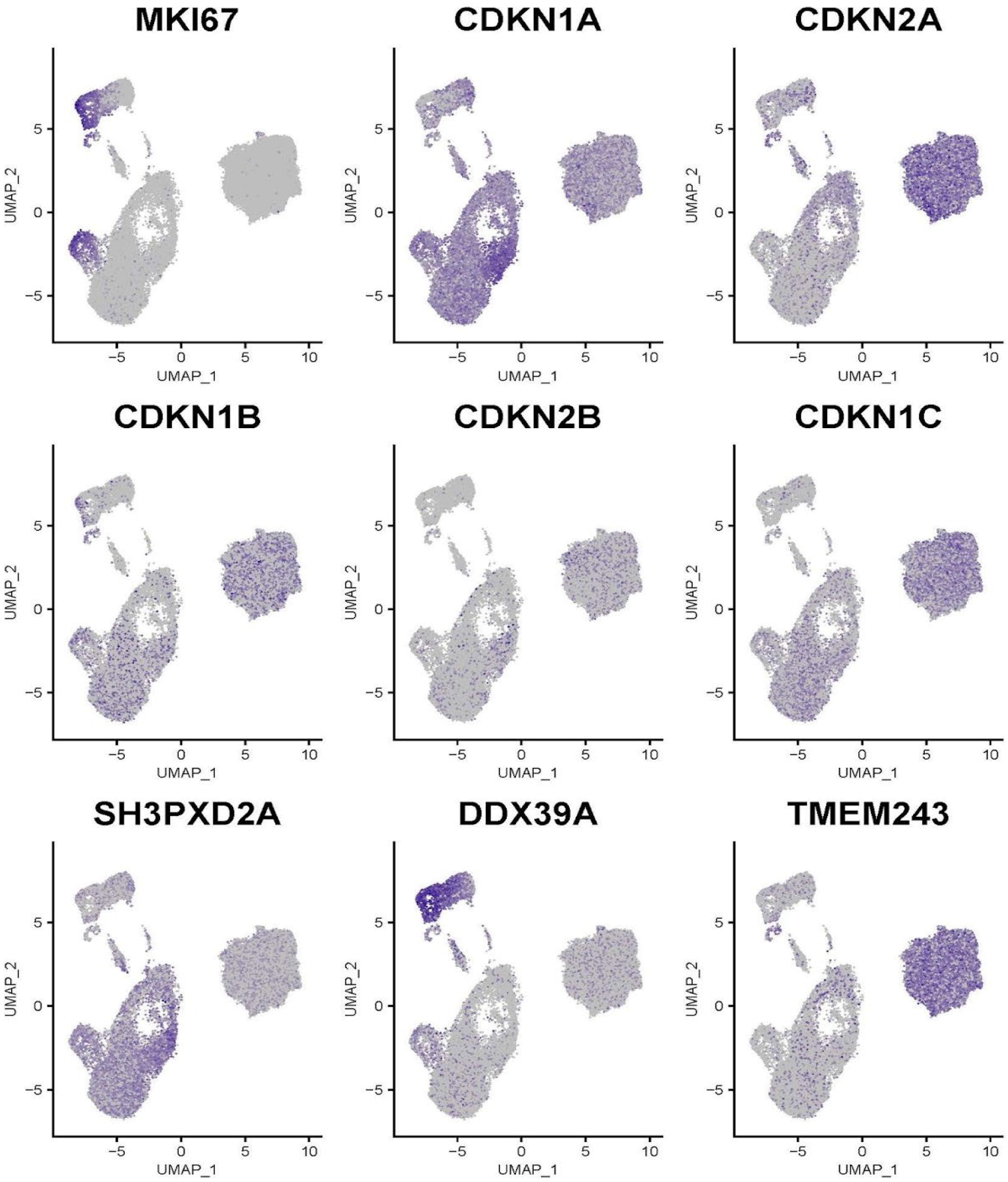
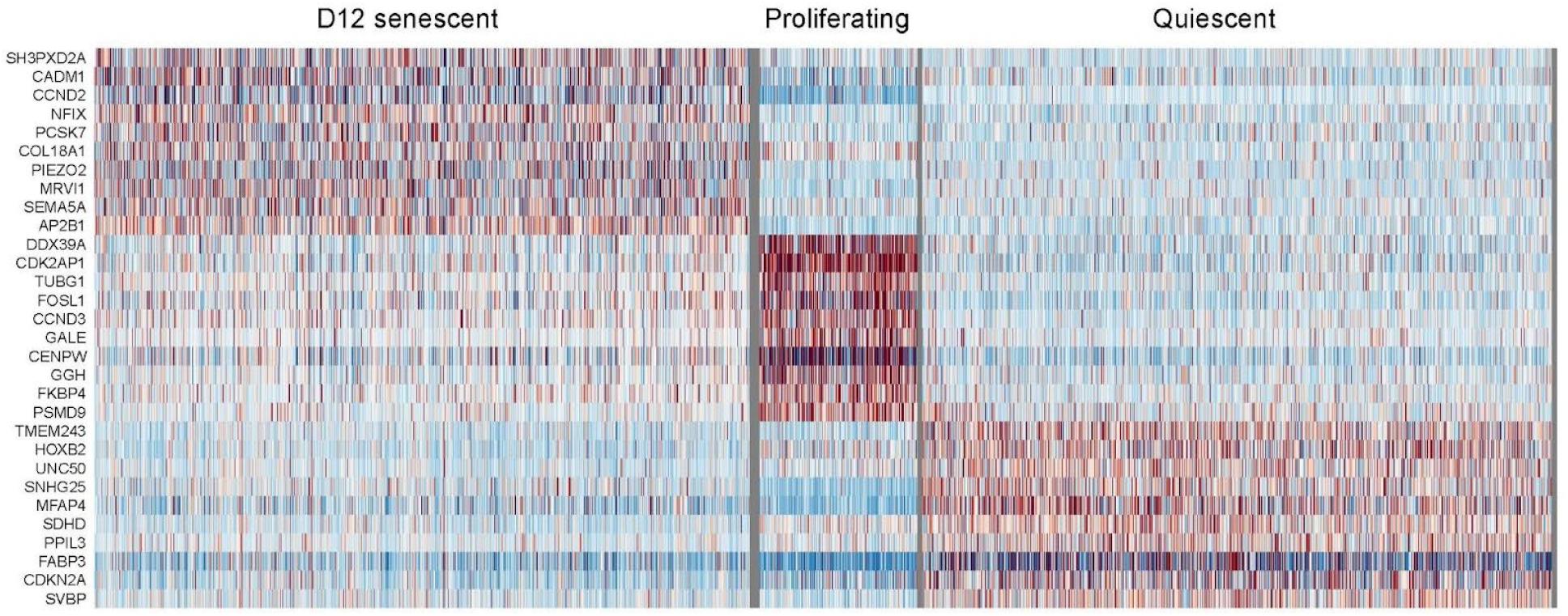
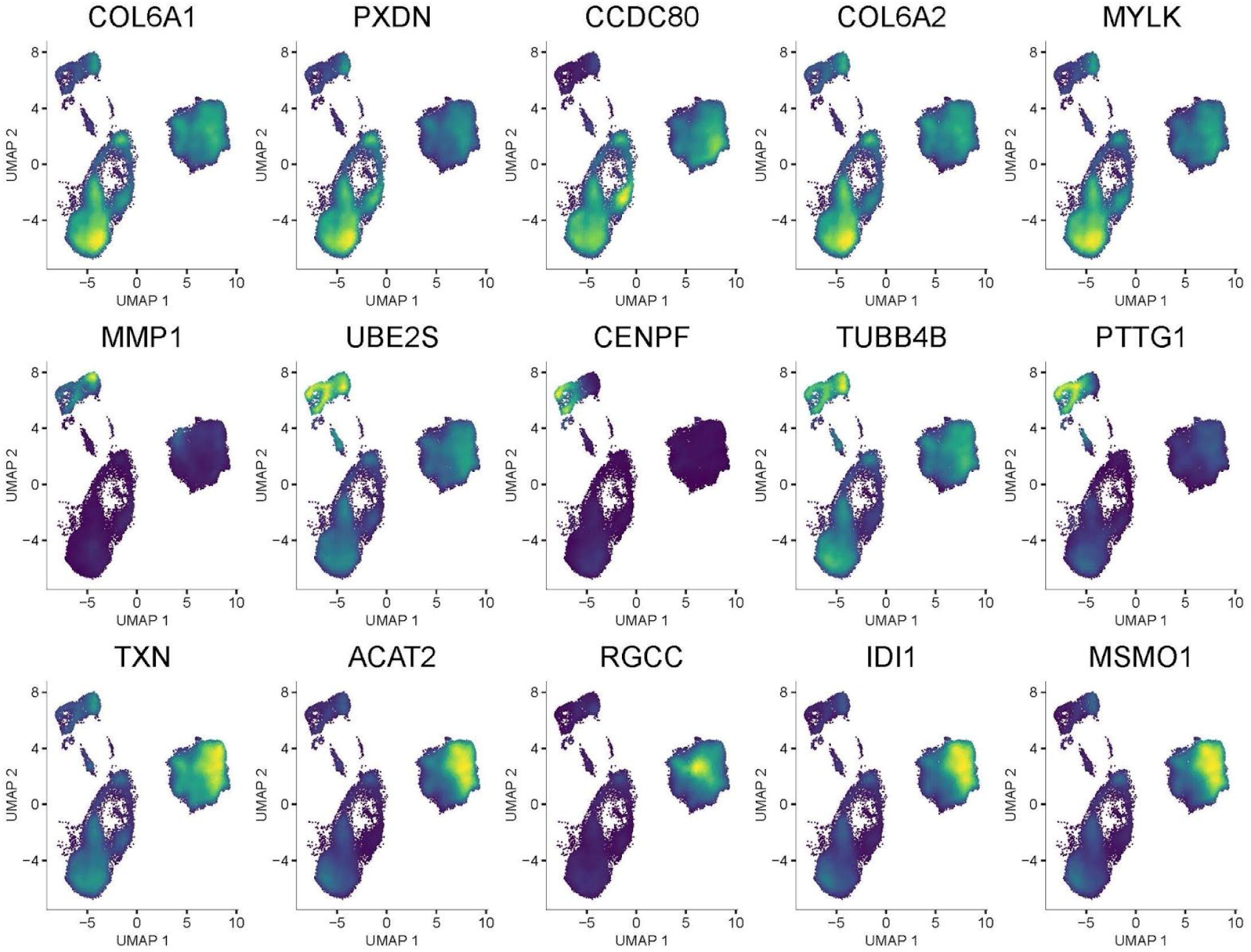
Cyclin-dependent kinase inhibitors are variably expressed in proliferating, quiescent, and senescent cells, which form heterogeneous populations identified by non-traditional markers. (A) Visualization of populations identified in comparisons of proliferating, quiescent, and SnCs; by time point (top) and cluster (bottom). (B) Scaled expression values of MKI67, a number of CDKN family genes, and three genes (SH3PXD2A, DDX39A, TMEM243) determined to be specific markers of cell state. (C) Expression heatmaps of genes identified as being specifically expressed in respective cell states. (D) Density plot of differentially expressed genes upregulated in senescent (top row), proliferating (middle), and QnCs (bottom).

The upregulation of cyclin-dependent kinase inhibitors CDKN1A (p21) and CDKN2A (p16 transcript) are frequently reported to be canonical markers of cell senescence. We compared the expression of these two genes, and their inhibitors CDKN1B (p27/Kip1), CDKN2B (p15/INK4b), and CDKN1C (P57/Kip2), in our cell states of interest. In SnCs, the most highly upregulated (by log fold change) of these two genes was CDKN1A, and was expressed in 92% of these cells. CDKN1A was also expressed in 71% of QnCs and 64% of proliferating cells, albeit to lower degrees. To our surprise, CDKN2A was most highly upregulated in QnCs (81%). Interestingly, 43% of proliferating cells also expressed CDKN2A, while only 28% of SnCs had significant expression. CDKN1B was downregulated in proliferating cells compared to quiescent and senescent cells, with the latter two cell states showing similar levels of expression. In small numbers of quiescent and proliferating cells, CDKN2B was slightly up- and downregulated respectively. CDKN1C was upregulated in both QnCs (67% of cells expressing) and SnCs (39%) compared to proliferating cells (13%). However, we did not measure protein levels for these genes at the level of the single cell.

The most specific gene expression marker in SnCs was SH3 and PX domain-containing protein SH3PXD2A (expressed in 66% of SnCs and only 33% of the other cells. RNA helicase DDX39A was both upregulated and a specific marker for proliferating cells (81%, compared to 20% of proliferating and QnCs). Transmembrane protein TMEM243 was found to be highly specific to QnCs, with 80% of them expressing the gene compared to only 17% of the remaining cells; other markers of considerable specificity for each cell state are plotted in Figure 3C. In this 3 way comparison, SnCs upregulated type VI collagen alpha chains COL6A1 and COL6A2, the ECM-associated peroxidase PXDN, secreted ECM component CCDC80, and myosin light chain kinase MYLK (Figure 3D). Upregulated genes in proliferating cells included matrix metallopeptidase MMP1, ubiquitin ligase UBE2S, centromere and microtubule protein CENPF, microtubule component TUBB4B, and pituitary tumor transforming PTTG1. Finally, QnCs upregulated thioredoxin TXN, acetyltransferase ACAT2, cell cycle regulator RGCC, peroxisomal enzyme IDI1, and monooxygenase MSMO1.

### Single cell transcriptomic time-course of senescence

In order to assess the full extent of heterogeneity within time points and across the time course of transformation to SnCs, we performed a merged analysis of proliferating (“Pro”) and post-doxo treatment (H12, D1, D2,…D12) samples. We adjusted clustering parameters in this analysis of 126,163 cells such that the population of senescence evaders (~6.5% of all cells) was restricted to one cluster. Figure 4A groups cells by time point, and the UMAP projection infers an unsynchronized progression through senescence. Apart from co-clustering of D1 cells, we identified a complex pattern of co-localization and overlap in cells from different timepoints. We found 46 (Figure 4B) distinct groups of cells, including the presence of proliferating cells from post-doxo treatment timepoints (Cluster 33).

**Figure 4:**
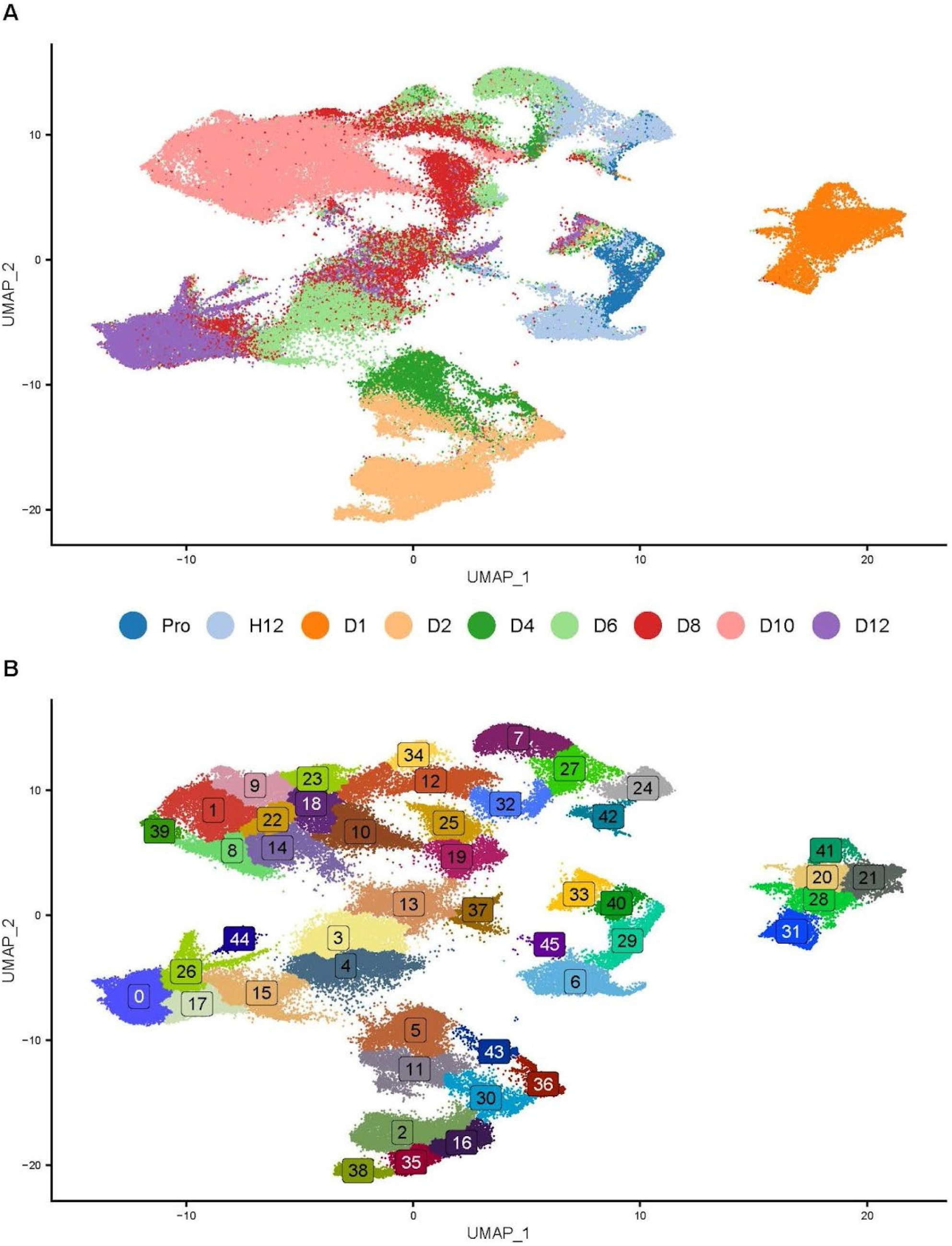
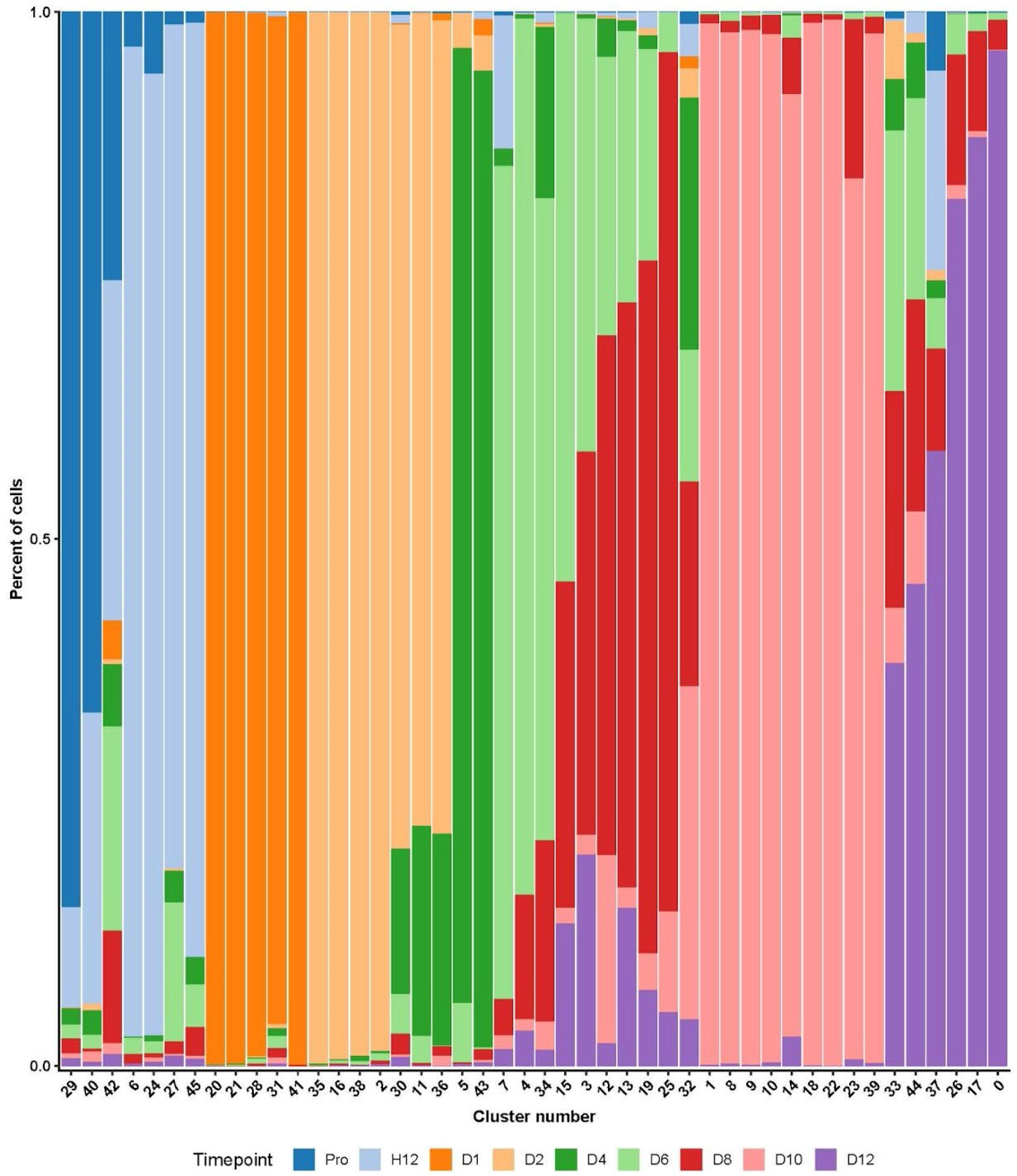
Visualization of senescence time-course. A) Clustering of 126,163 cells across 9 time points, plotted with the UMAP dimensional reduction technique. Pro = proliferating cells, H12 = 12 hours after doxo treatment, D1 = 1 day after doxo treatment, D2 = 2 days after doxo treatment, and so on up to D12. B) Louvain clustering of merged data identifies 46 distinct clusters across the full time-course. C) Cluster composition plotted by timepoint.

Clusters were composed of non-homogenous populations of cells from multiple time points (Figure 4C). Clusters 0, 17, 26, 33, 37, and 44 are dominated by cells from D12, suggesting these are clusters of stable SnCs. However, many cells from earlier timepoints also co-cluster with these D12 cells. This suggests that with senescence induction, some cells quickly (as soon as 12 hours post doxo treatment) reorient their transcriptional profiles from a proliferative potential to senescent. Conversely, some cells appear to lag in their transformation to SnCs. This is illustrated by the presence of D12 cells in clusters dominated by cells from timepoints D4/D6/D8 (Clusters 3, 4, 12, 13, 15, 19, 25, 32, 34). Our study shows that the development of senescence is a dynamic process, through which cells do not progress in lock-step. This emphasizes the value of single cell profiling in the senescent paradigm, while raising concerns about bulk preparation workflows of SnCs of either protein or gene expression. Evaluating the progression of senescence across the experimental timeline provided insight into the heterogeneity among cell populations at each timepoint. However, this approach did not facilitate identifying and tracking cell trajectories across sequential timepoints. To capture greater transcriptional specificity in smaller timescales, we carried out integrated analyses of subsets of the time course.

### Validating markers of proliferation and senescence across transcriptomic time course

In order to validate cell states over the full time course, we examined the expression patterns of known markers of proliferation and senescence. We focused on MKI67, TOP2A, and CCNA2 due to their widely recognized roles as markers of cycling cells. These three genes were expressed in untreated proliferating cells and are downregulated as quickly as 12 hours after senescence induction. Furthermore, they are sporadically expressed in subpopulations of cells from post-doxo treatment time points (Figure 5A).

**Figure 5:**
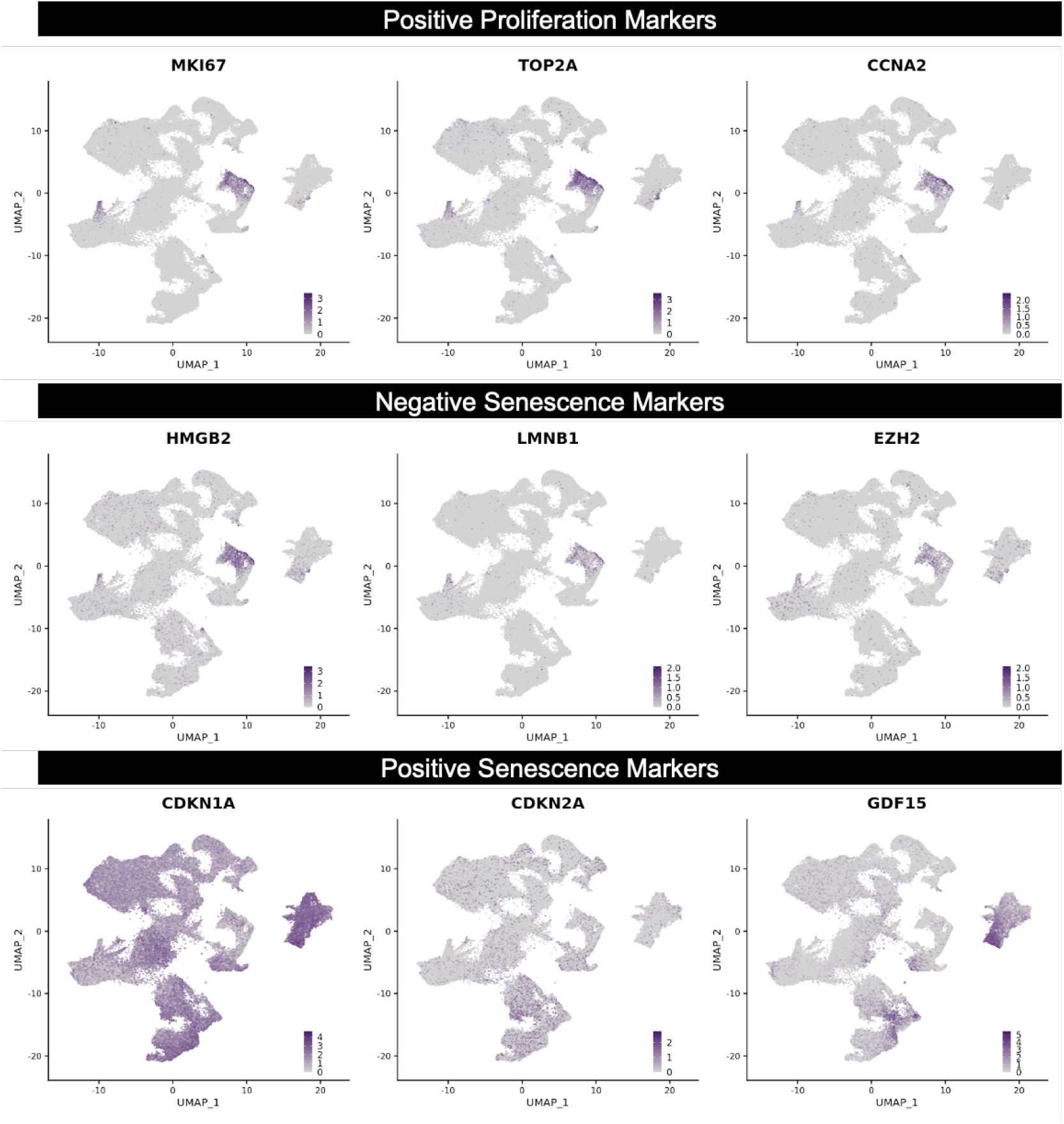
Normalized expression of markers of proliferation and senescence. Darker purple shading indicates increased expression. A) Genes MKI67 (left), TOP2A (middle), and CCNA2 (right) were predominantly expressed in clusters of proliferating cells. B) Genes HMGB2 (left), LMNB1 (middle), EZH2 (right), all negative senescence markers, were also expressed in the clusters of proliferating cells. C) CDKN1A (left), CDKN2A (middle), and GDF15 (right), all positive markers of senescence, were upregulated to different degrees following doxo treatment.

We also examined expression patterns of negative markers for proliferation HMGB2, LMNB1, and EZH2 (Figure 5B), genes whose proteins are reported to be downregulated as cells enter senescence. Interestingly, these markers are not often detected via common bulk RNA analyses, and changes in their expression at the RNA level have been deliberated in the senescence field. Here, we demonstrate that transcriptional changes of these markers can be captured using a time course based single cell approach. We investigated how the expression of the established positive senescence markers CDKN1A, CDKN2A, and GDF15 change across the time course we employed in this study. The expression of CDKN1A/p21 was widely distributed - its low, sporadic presence in proliferating cells may be due to its role in transitory cell cycle arrest at the G1 checkpoint. As early as 12 hours after senescence induction we observed a transient increase in its gene expression, which then increased from D1 to D4. By D12, its expression was lower, which may be attributed to cells reaching stable arrest, no longer necessitating p21 overproduction. In contrast to CDKN1A, CDKN2A (p16 and p14arf) expression was much weaker throughout the time course, despite an increase at timepoints D2 and D4 when compared to proliferating cells. The cytokine GDF15 is an important senescence associated secretory phenotype (SASP) molecule, whose secretion plays a role in apoptosis and inflammation signaling. We observed an early upregulation of GDF15 from 12-24 hours (H12 and D1), before subsiding starting at D2.This finding contrasts with the unchanged CDKN2A levels from H12 to D1. Altogether, these findings serve as confirmation of senescence induction using several conventional senescence markers.

### Identification of cell trajectories across early, middle, and late phases of senescence

We next investigated whether distinct lineages of senescent cells could be identified. We separated the total time course into three distinct phases: early (proliferating to D2), middle (D2 to D8), and late-stage senescence (D8 to D12), allowing us to explore each with greater specificity. These temporal stages were chosen in conjunction with normalization of expression levels of CDKN1A (Figure S3), as well as changes in ambient RNA fraction (Figure 1). Following processing by the Seurat pipeline, the Slingshot and tradeSeq R packages were utilized to perform pseudotime analysis and visualize the transcriptional shifts along identified trajectories. These analysis packages were chosen due to their ability to identify multiple trajectories in heterogeneous single cell data. Of particular interest were clusters in which a given lineage diverged from other lineages, and clusters in which the given lineage ended (putatively the most temporally advanced cells of a lineage). The top up- and down-regulated genes (by log2 fold change) were identified for the divergence and end clusters, as well as the most specific up and down-regulated genes (by percent cells expressing a given gene). Early stage senescence is composed of H12, D1, and D2 cells. All early senescence trajectories commenced in cluster 18, largely composed of proliferating cells, before splitting off into one of six lineages (Figure 6a).

**Figure 6:**
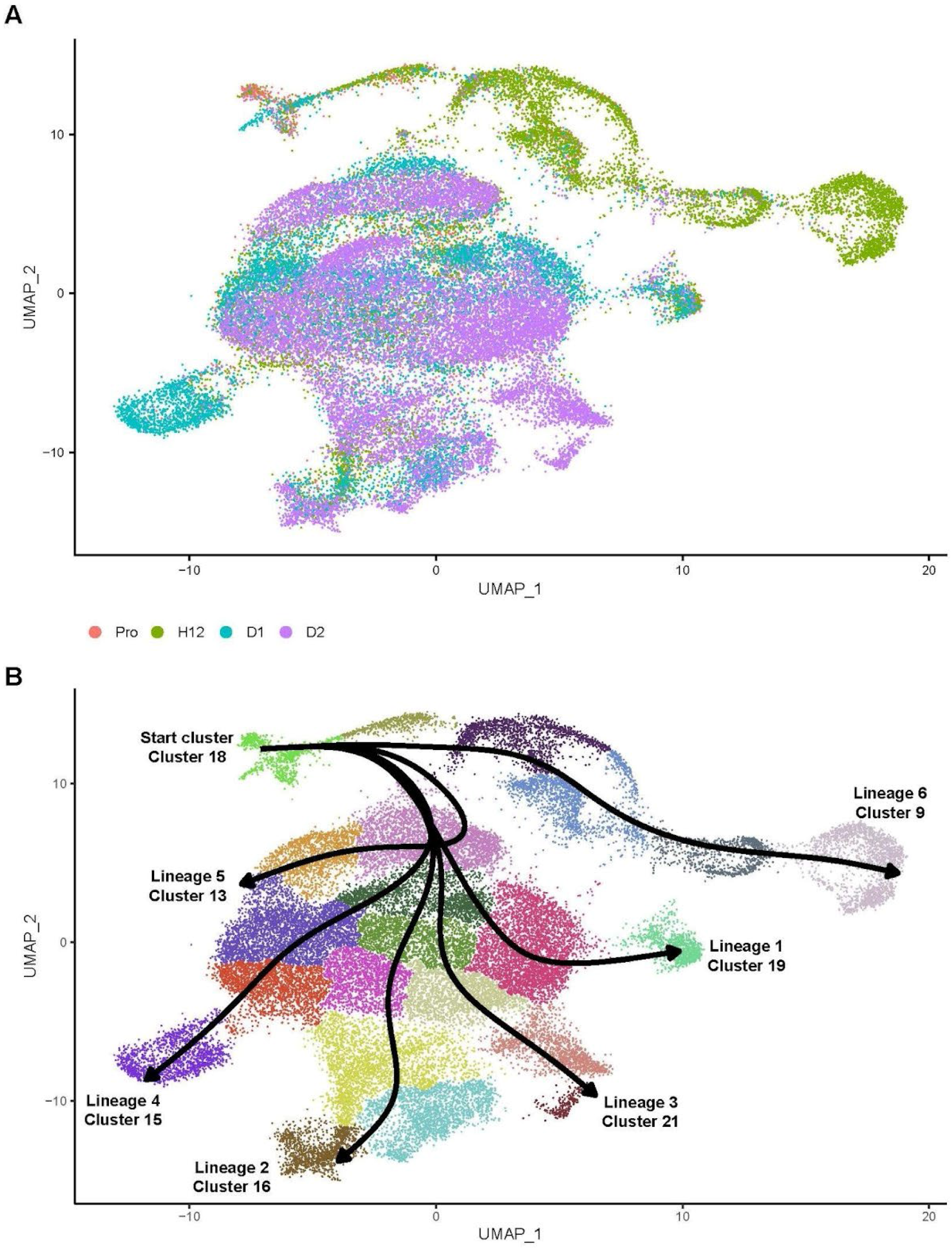
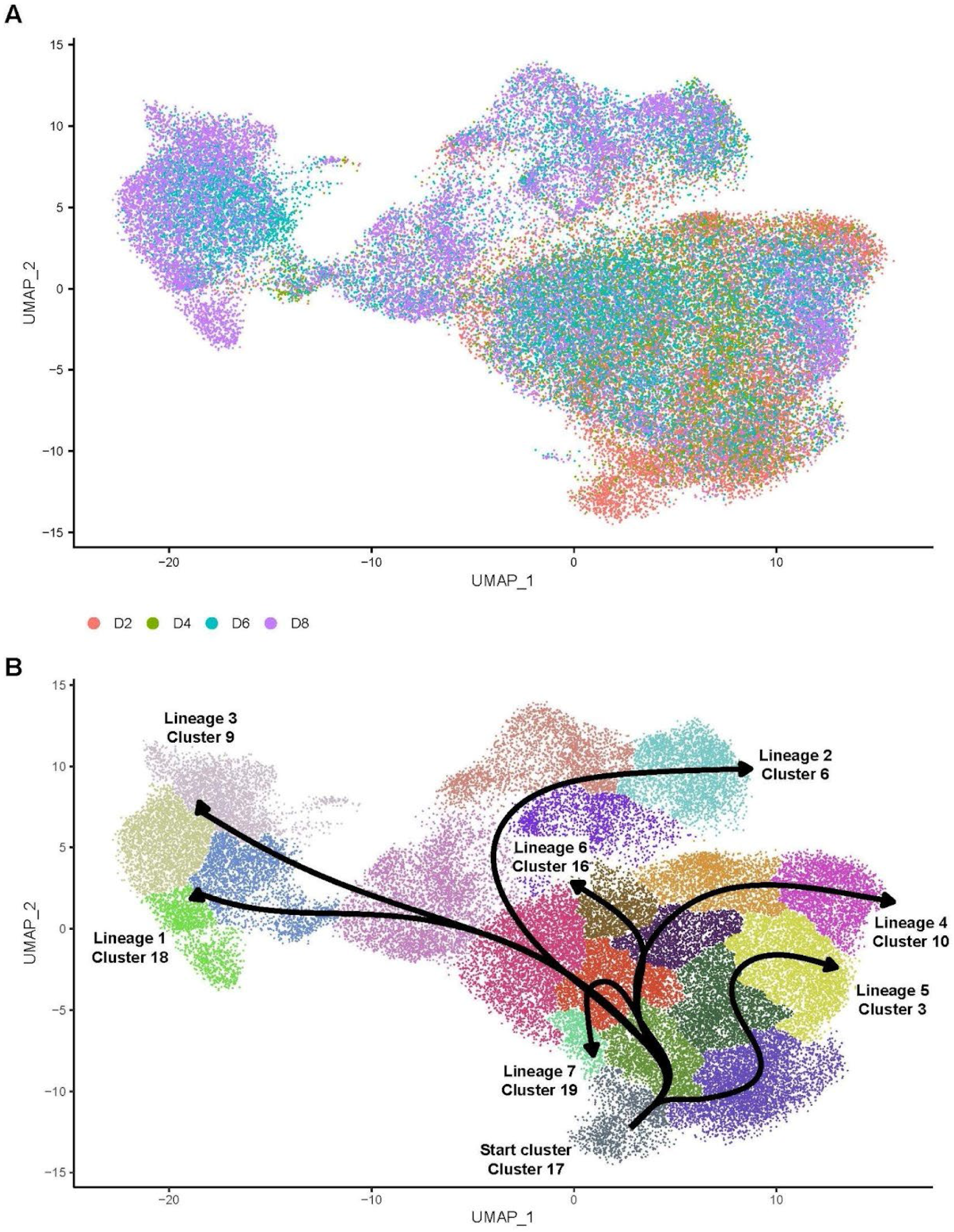
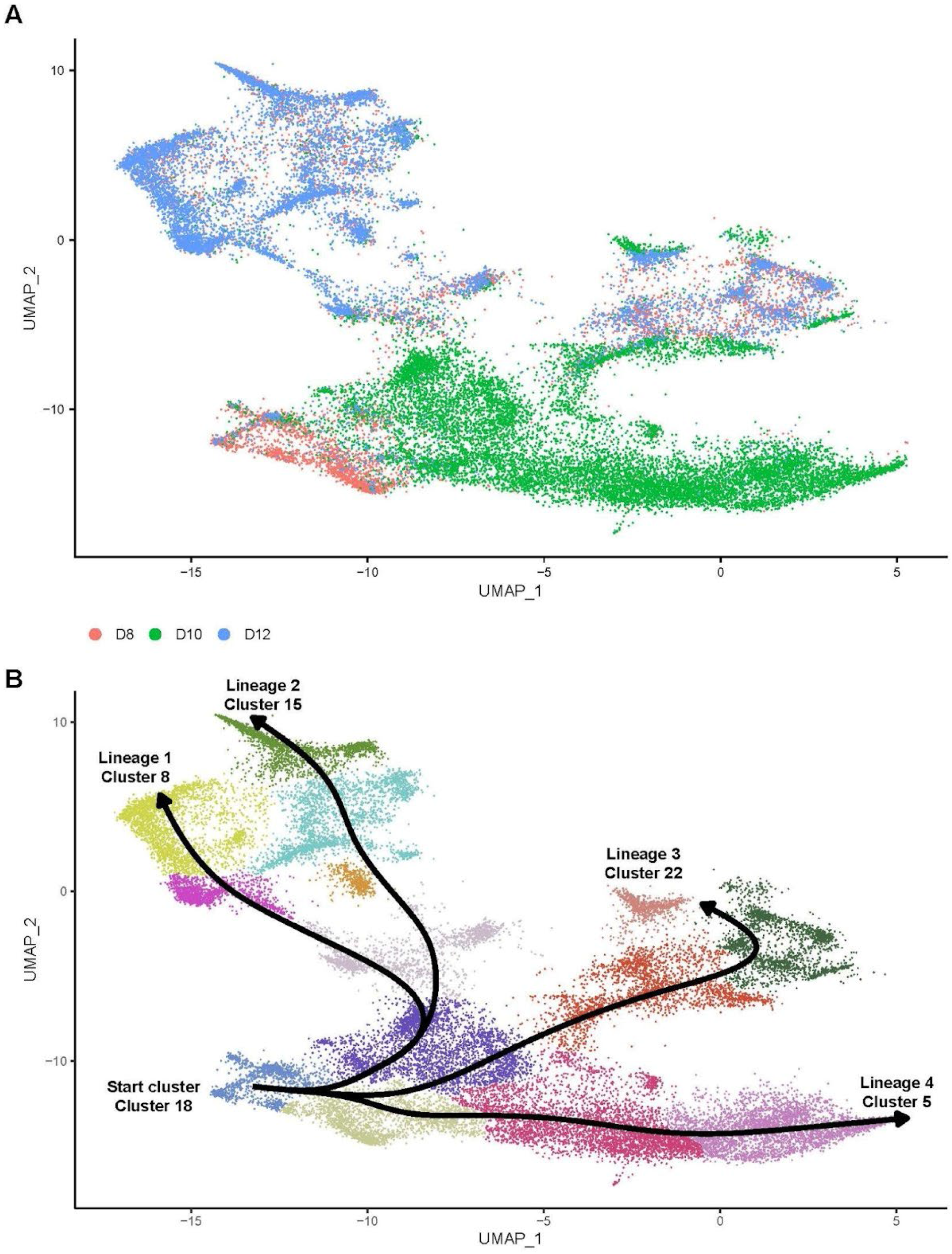
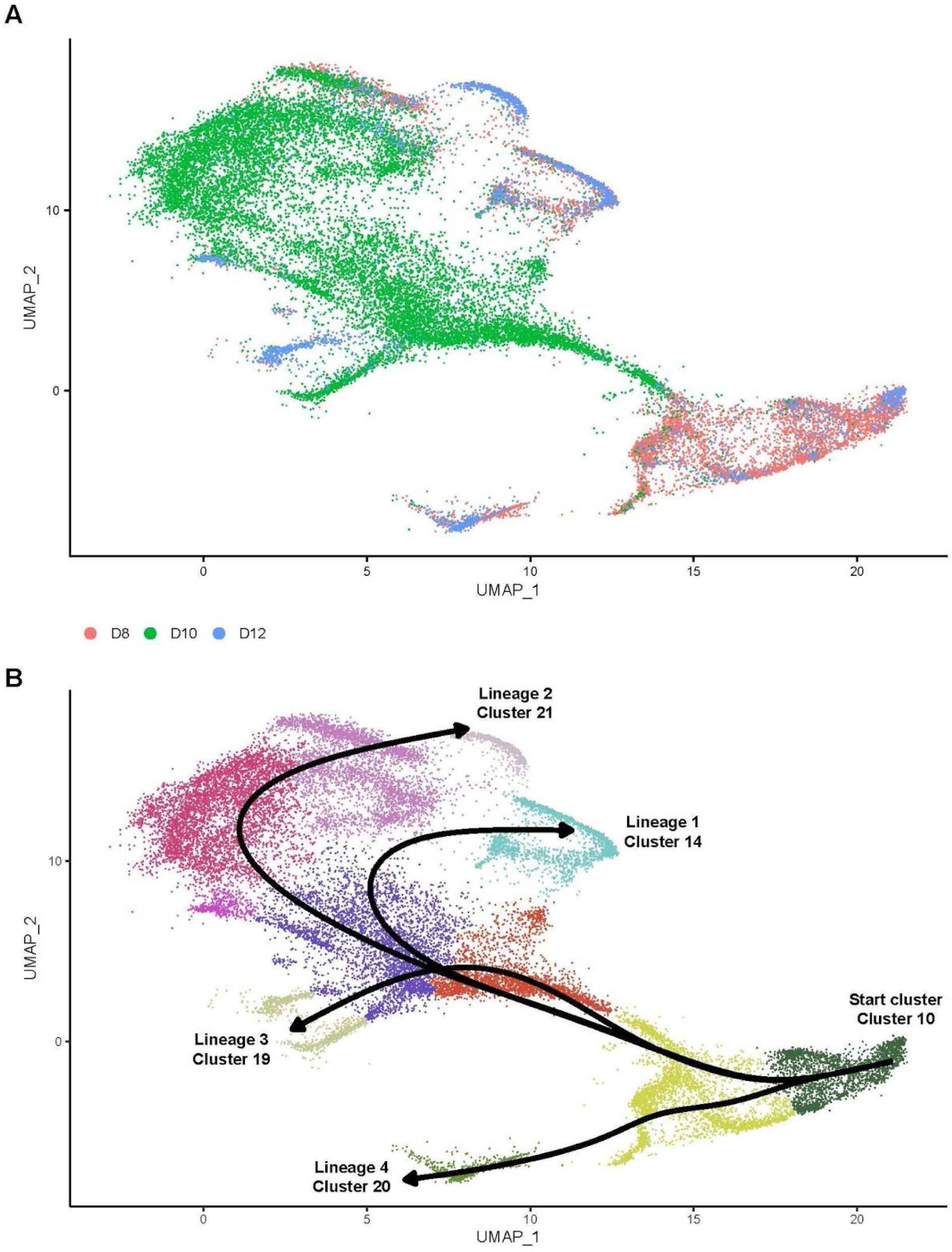
Senescing cells form distinct transcriptional lineages as they progress from proliferation through late-stage senescence. Lineage trajectories are identified by number and end cluster. (A) From untreated proliferating cells to the D2 timepoint, early stage senescence trajectories begin at cluster 18 and mature down one of six lineages, ending in clusters 19, 16, 21, 15, 13, and 9. (B) Mid-stage senescence (time points D2 to D8) diverges into seven lineages which start at cluster 17 and end in clusters 18, 6, 9, 10, 3, 16, and 19. (C) Late-stage senescence (D8 - D12) lineages originated at clusters 18 and 10. Four lineages start from both cluster 18 (lineages ending in clusters 8, 15, 22, 5) and (D) cluster 10 (lineages ending in clusters 14, 21, 19, and 20).

**Figure 7:**
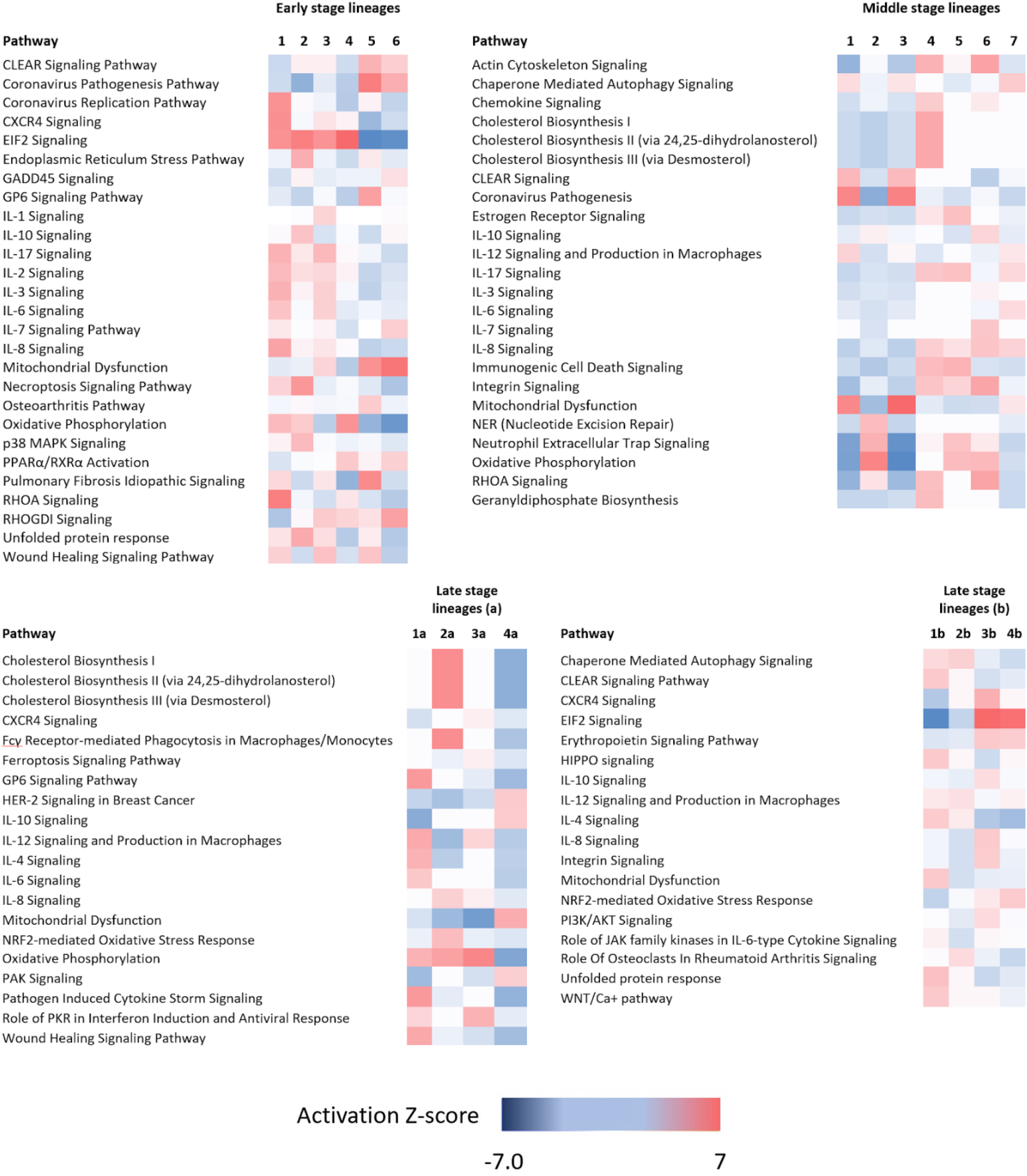
Differentially regulated pathways in senescence lineages. Pathway activation z-scores were calculated in IPA. Pathway activation/inhibition are denoted by positive/negative z-scores. **(A)** Early stage senescence is characterized by lineages that differentially express stress response, inflammatory, and DNA damage related pathways. **(B)** Pathway activity in middle stage senescence lineages. **(C) - (D)** Pathway activity in two branches of late stage senescence lineages.

For each lineage, we identified genes with positive expression and specificity to their cluster relative to others (Table 3), as well as negative changes (Table 4). Positive changes within early-stage senescence were genes associated with response to oxidative stress (TP53I3, lineage 1) and inflammatory response (CXCL8, lineage 3).

**Table 3.** Upregulated (UR) lineage markers of early-stage senescence (Pro-D2). Top genes were determined by highest Log2 fold change. Specificity was determined by change in the percent of cells expressing a gene of interest. For both divergence and end clusters, comparisons were performed by comparing a given cluster against all other clusters of that type.

| Lineage | Lineage Divergence Cluster | Top UR log2FC gene | Top UR gene by specificity | Lineage End Cluster | Top UR log2FC gene | Top UR gene by specificity |
| --- | --- | --- | --- | --- | --- | --- |
| Lineage 1 | 0 | TP53I3 | ARHGAP22 | 13 | ACAT2 | TUBB2A |
| Lineage 2 | 3 | C12orf75 | IGFBP3 | 15 | GDF15 | PSAT1 |
| Lineage 3 | 12 | CXCL8 | AREG | 21 | CXCL8 | FOS |
| Lineage 4 | 1 | IGFBP3 | IGFBP3 | 16 | BEX3 | TP53TG1 |
| Lineage 5 | 13 | CTGF | CHIC2 | 19 | CD63 | EMILIN1 |
| Lineage 6 | 17 | MYL9 | ACTG2 | 9 | NABP1 | CPNE7 |

**Table 4.** Downregulated (DR) lineage markers of early-stage senescence (Pro-D2). Top Log2FC genes were determined by highest Log2 fold change, while specificity was determined by change in the percent of cells expressing a given gene.

| Lineage | Lineage Divergence Cluster | Top DR log2FC gene | Top DR gene by specificity | Lineage End Cluster | Top DR log2FC gene | Top DR gene by specificity |
| --- | --- | --- | --- | --- | --- | --- |
| Lineage 1 | 0 | CXCL8 | AREG | 13 | GDF15 | POLR2J3 |
| Lineage 2 | 3 | CLU | LRPAP1 | 15 | IGFBP5 | IGFBP5 |
| Lineage 3 | 12 | TP53I3 | ARHGAP22 | 21 | NABP1 | CDC42EP3 |
| Lineage 4 | 1 | COL3A1 | SOX4 | 16 | GDF15 | DDX5 |
| Lineage 5 | 13 | MMP1 | MMP1 | 19 | TXN | BLOC1S2 |
| Lineage 6 | 17 | SERPINE2 | EMC7 | 9 | GDF15 | EMC7 |

Lineage 1 diverged from the rest at cluster 0, which upregulated the p53 inducible oxidoreductase TP53I3 and the GTPase activator ARHGAP22 (Table 3). This lineage ended at cluster 13, marked by expression of acetyltransferase ACAT2 and tubulin gene TUBB2A. Expression of C12orf75 and IGFBP3 in cluster 3 marked the divergence of lineage 2, which ended in cluster 15, which expresses growth factor GDF15 and aminotransferase PSAT1. SASP factor CXCL8 was also upregulated across lineage 3, from its divergence point (cluster 12) until its end cluster 21, in which the AP-1 transcription factor component FOS is expressed. The divergence of lineage 4 occurred in cluster 1, where IGFBP3 was both the most highly upregulated gene and most specific marker for this cluster. This lineage ended in cluster 16, which expressed BEX3, a protein involved in p75NTR-mediated apoptosis. Interestingly, the most specific marker for this cluster was TP53TG1, a lncRNA with both pro- and anti-oncogenic properties. Lineage 5 expressed CTGF and CHIC2 when diverging, before ending in cluster 19 which expressed cell surface protein CD63 and extracellular matrix glycoprotein EMILIN1. The final early-stage lineage was number 6, which began its trajectory by expressing myosin chain MYL9 and smooth muscle actin expressing ACTG2. This lineage ended in cluster 9, concomitant with upregulation. of NABP1 and CPNE7. Interestingly, SASP genes including AREG, CLU, CXCL8, MMP1, and GDF15 were downregulated in different early-stage lineages (Table 4). Our data shows that early in senescence, cells separate into lineages with distinct transcriptional profiles and specific expression of both SASP and non-SASP genes. Differential regulation of pathways provided insight into functional specializations of some SnC lineages.

We determined that in lineage 1, CXCR4, IL-8, actin cytoskeleton (RHOA, Rho GTPase, and RAC), and ephrin receptor signaling were all activated. The coronavirus replication pathway was also uniquely activated in lineage 1, while the virus’ pathogenesis pathway was expressed in lineages 5 and 6 only. Lineage 2 was marked by increased endoplasmic reticulum stress signaling, p38/MAPK activity, and inflammatory signaling mediated by IL-10. The unfolded protein response (UPR) activity was highest in this lineage, as was necroptosis signaling. The population of developing SnCs in lineage 3 was characterized by IL-17 and hypercytokinemia signaling. Lineage 4 modulates a number of pathways shared with other lineages, such as EIF2 signaling (also activated in lineages 1-3, inhibited in lineages 5 and 6), oxidative phosphorylation (shared with lineages 1 and 2, inhibited in others) and PPARα/RXRα activation (shared only with lineage 6). However, no strongly specific or uniquely activated pathways could be determined. Pathways activated in lineage 5 include osteoarthritis, collagen receptor GP6, and pulmonary fibrosis signaling. The mitochondrial dysfunction pathway was activated in lineages 3, 5, and 6. The latter two lineages also displayed increased lysosomal activity (CLEAR pathway). Wound healing, one of the beneficial roles SnCs play, was activated in lineages 1, 3, and 5. Finally, in lineage 6, GADD45 (Growth Arrest and DNA Damage-inducible 45 family genes) signaling was uniquely upregulated.

Middle stage senescence was composed of D2, D4, D6, and D8 cells, which were clustered and visualized separately from other timepoints or stages of senescence. The UMAP projection shows a gradual progression of cells from Day 2 to Day 8 after senescence induction. Cells started their trajectories in cluster 17 (Figure 6b), largely made up of D2 cells, before proceeding into one of seven lineages. As with early senescence lineages, we observed a large degree of heterogeneity in this sequence of time points. While the end clusters of lineages 1 (cluster 18), 2 (cluster 6), 3 (cluster 9), and 5 (cluster 3) are composed mainly of cells from D8, non-terminal clusters had mixed populations of cells from all timepoints. For each lineage, we identified genes with positive expression changes and positive specificity to their cluster relative to the others (Table 5), as well as negative changes (Table 6).

**Table 5.**
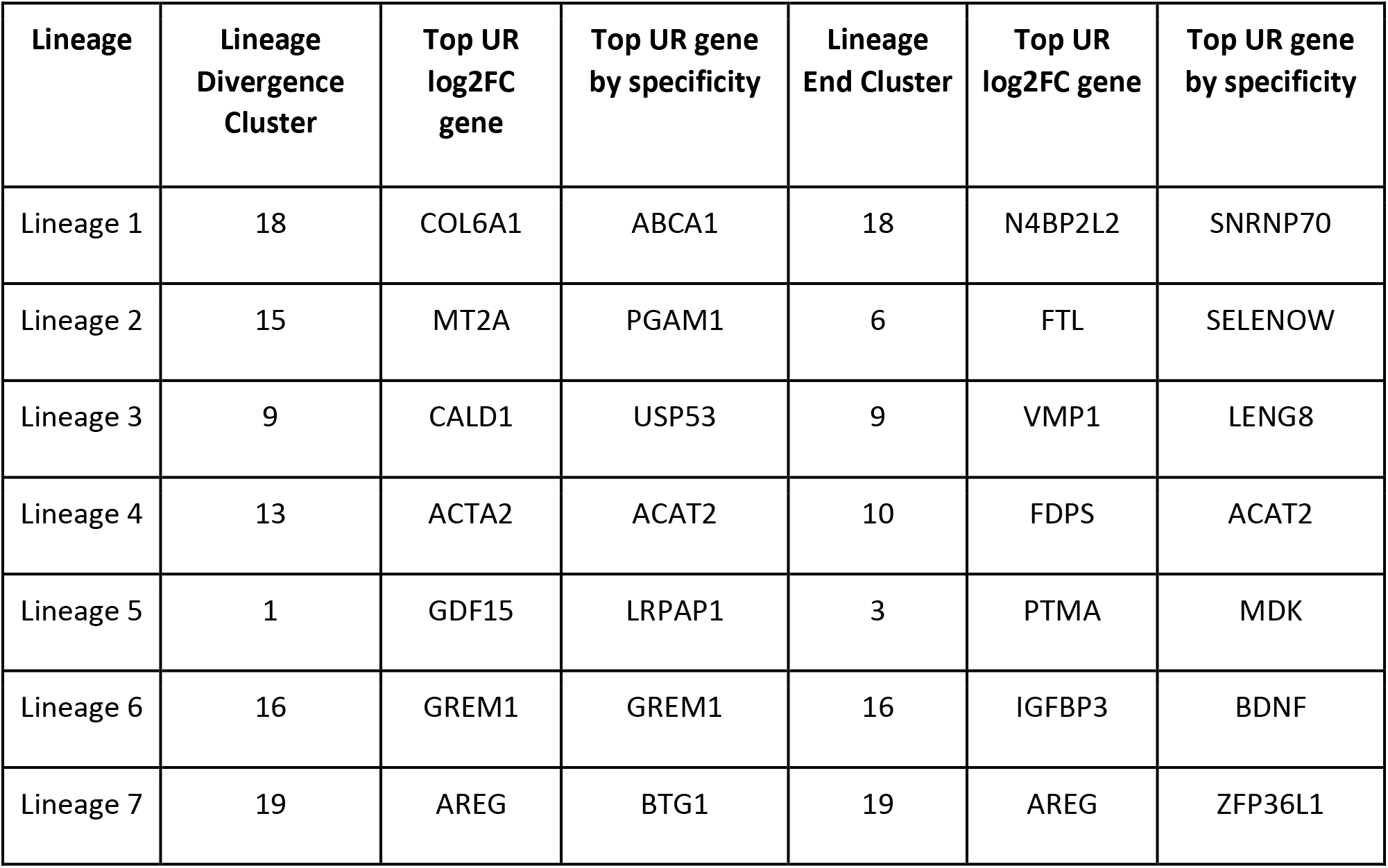
Upregulated (UR) lineage markers of mid-stage senescence (D2-8). Top Log2FC genes were determined by highest Log2 fold change, while specificity was determined by change in the percent of cells expressing a given gene.

| Lineage | Lineage Divergence Cluster | Top UR log2FC gene | Top UR gene by specificity | Lineage End Cluster | Top UR log2FC gene | Top UR gene by specificity |
| --- | --- | --- | --- | --- | --- | --- |
| Lineage 1 | 18 | COL6A1 | ABCA1 | 18 | N4BP2L2 | SNRNP70 |
| Lineage 2 | 15 | MT2A | PGAM1 | 6 | FTL | SELENOW |
| Lineage 3 | 9 | CALD1 | USP53 | 9 | VMP1 | LENG8 |
| Lineage 4 | 13 | ACTA2 | ACAT2 | 10 | FDPS | ACAT2 |
| Lineage 5 | 1 | GDF15 | LRPAP1 | 3 | PTMA | MDK |
| Lineage 6 | 16 | GREM1 | GREM1 | 16 | IGFBP3 | BDNF |
| Lineage 7 | 19 | AREG | BTG1 | 19 | AREG | ZFP36L1 |

**Table 6.** Downregulated (DR) lineage markers of mid-stage senescence (D2-D8). Top Log2FC genes were determined by highest Log2 fold change, while specificity was determined by change in the percent of cells expressing a given gene.

| Lineage | Lineage Divergence Cluster | Top DR log2FC gene | Top DR gene by specificity | Lineage End Cluster | Top DR log2FC gene | Top DR gene by specificity |
| --- | --- | --- | --- | --- | --- | --- |
| Lineage 1 | 18 | CALD1 | USP53 | 18 | TXN | DBI |
| Lineage 2 | 15 | DST | N4BP2L2 | 6 | COL1A1 | CYR61 |
| Lineage 3 | 9 | COL6A1 | ABCA1 | 9 | TXN | SUMO2 |
| Lineage 4 | 13 | GREM1 | GREM1 | 10 | VMP1 | SELENOW |
| Lineage 5 | 1 | C12orf75 | CDC42EP3 | 3 | CCDC80 | CCDC80 |
| Lineage 6 | 16 | ACTA2 | ACAT2 | 16 | VMP1 | TKT |
| Lineage 7 | 19 | IGFBP3 | MYL12A | 19 | PRDX1 | MYL12A |

Mid stage lineage 1 expressed the collagen gene COL6A1, the ABCA1 cholesterol phospholipid transporter, transcription repressor complex component N4BP2L2, and small nuclear ribonucleoprotein SNRNP70. Metallothionein MT2A, glycolytic mutase PGAM1, the iron storage ferritin chain FTL, and glutathione dependent antioxidant SELENOW were upregulated over lineage 2. Actin-binding protein CALD1, thioredoxin TXN, ubiquitin-like modifier SUMO2, and ubiquitin peptidase USP53 were upregulated by cells in cluster 18 / lineage 3. Lineage 4 was defined by upregulation of smooth muscle actin ACTA2, acetyltransferase ACAT2, and farnesyl diphosphate synthase FDPS. GDF15 was again upregulated in lineage 5, along with LDL receptor chaperone LRPAP1, antiapoptotic protein PTMA, and secreted growth factor MDK. Bone morphogenic protein antagonist GREM1 and nerve growth factor BDNF were the strongest markers of lineage 6. Finally, AREG, anti-proliferative factor BTG1, and zinc finger-like protein ZFP36L1 were upregulated in lineage 7.

Lineage 1 and 3 shared many activated pathways, most significantly the coronavirus pathogenesis pathway and mitochondrial dysfunction, as well as CLEAR (Coordinated Lysosomal Expression and Regulation). IL-12 signaling was activated in lineages 1 and 6. Oxidative phosphorylation and neutrophil extracellular trap signaling were upregulated in lineages 2, 5, and 6. Nucleotide excision repair was uniquely activated in lineage 2. Lineage 4 was unique in its activation of numerous cholesterol synthesis pathways, as well as geranylgeranyl diphosphate biosynthesis. Lineages 4 and 5 shared many activated pathways, including estrogen receptor and immunogenic cell death signaling.

Despite differences at the DEG level, the number of shared pathways between the two lineages indicate that some cells may take different paths to the same cell fate. Actin cytoskeleton, RHOA, and integrin signaling were all activated in lineage 6, indicating a cell lineage undergoing cytoskeletal reorganization and cell surface remodeling. Il-10 signaling was upregulated in lineages 6 and 2, and lastly IL-6 signaling was uniquely activated in lineage 7. IL-8 was activated in lineages 4-7 but inhibited in lineages 1-3. From D2 to D8, SnCs evolve into lineages characterized by signaling by specific SASP cytokines, metabolic activity, and cytoskeletal remodeling. Wound healing signaling was activated in lineages 4-7, continuing a trend identified in the early stage lineages.

Late-stage senescence was defined by presence of D8, D10, and D12 cells. In contrast to early and middle stages, these cells initiated their lineage start in one of two distinct clusters highly enriched for D8 cells (add percentage of composition?). The first subset of cells (a) began their trajectories in cluster 18 (Figure 6c) before splitting off into one of four lineages (lineages 1-4a ending in clusters 8, 15, 22, and 5 respectively). The second subset also evolved into 4 lineages after starting in cluster 10 (Figure 6d); these were named lineages 1-4b, and ended in clusters 14, 21, 19, and 20. As with the analyses of previous stages, D8-D12 cells were embedded and clustered separately from other time points. Lineage 1a upregulated neuronal pentraxin NPTX1 (Table 7), retinol binder RBP1, COL3A1, and procollagen endopeptidase PCOLCE. Follistatin (FST), squalene epoxidase (SQLE), and again ACAT2 were markers for lineage 2a. Transcription factors KLF6 and SOX4 were upregulated in lineage 3a, alongside secreted Wnt signaling modulator SFRP1 and histone HIST1H4C. Lineage 4 upregulated myofibroblast marker TPM2, antiapoptotic lncRNA MTRNR2l12, and IGFBP7. In the second subset of late-stage senescence trajectories, lineage 1b expressed the metalloproteinase and IGFBP-family cleaving PAPPA when diverging from other lineages. IGFBP3 and carboxypeptidase CPA4 were upregulated at the end of this lineage. Collagen chain COL6A3 and secreted growth factor PTN were upregulated in lineage 2b, across the entire trajectory from divergence until the end cluster (cluster 21). Lineage 3b upregulated phosphodiesterase PDE5A and the lncRNA NORAD, which is produced in response to DNA damage, when diverging at cluster 19. This lineage also expressed YBX1, a secreted cold shock domain protein that functions as an extracellular mitogen, and SET, a nuclear protein that inhibits apoptosis and histone acetylation. The final lineage, 4b, was characterized by strong and specific expression of GDF15, unique among the late state lineages.

**Table 7.** Upregulated (UR) lineage markers of late-stage senescence (D8-D12). Top Log2FC genes were determined by highest Log2 fold change, while specificity was determined by change in the percent of cells expressing a given gene.

| Lineage | Lineage Divergence Cluster | Top UR log2FC gene | Top UR gene by specificity | Lineage End Cluster | Top UR log2FC gene | Top UR gene by specificity |
| --- | --- | --- | --- | --- | --- | --- |
| Lineage 1a | 17 | NPTX1 | RBP1 | 8 | COL3A1 | PCOLCE |
| Lineage 2a | 15 | FST | SQLE | 15 | FST | ACAT2 |
| Lineage 3a | 9 | SFRP1 | KLF6 | 22 | SOX4 | HIST1H4C |
| Lineage 4a | 2 | TPM2 | MTRNR2L12 | 5 | IGFBP7 | IGFBP7 |
| Lineage 1b | 14 | IGFBP3 | PAPPA | 14 | IGFBP3 | CPA4 |
| Lineage 2b | 21 | COL6A3 | PTN | 21 | COL6A3 | PTN |
| Lineage 3b | 19 | PDE5A | NORAD | 19 | YBX1 | SET |
| Lineage 4b | 20 | GDF15 | GDF15 | 20 | GDF15 | GDF15 |

**Table 8.** Downregulated (DR) lineage markers of late-stage senescence (D8-D12). Top Log2FC genes were determined by highest Log2 fold change, while specificity was determined by change in the percent of cells expressing a given gene.

| Lineage | Lineage Divergence Cluster | Top DR log2FC gene | Top DR gene by specificity | Lineage End Cluster | Top DR log2FC gene | Top DR gene by specificity |
| --- | --- | --- | --- | --- | --- | --- |
| Lineage 1a | 17 | FST | SQLE | 8 | CCND1 | GAS6 |
| Lineage 2a | 15 | NPTX1 | RBP1 | 15 | TIMP1 | IGFBP7 |
| Lineage 3a | 9 | TIMP1 | PRDX1 | 22 | PRDX1 | ANXA1 |
| Lineage 4a | 2 | FOSB | FOSB | 5 | FOSB | FOSB |
| Lineage 1b | 14 | COL6A3 | PTN | 14 | PTMA | PTMA |
| Lineage 2b | 21 | IGFBP3 | PAPPA | 21 | IGFBP3 | PAPPA |
| Lineage 3b | 19 | TUBA1B | TUBB4B | 19 | TIMP1 | LRPAP1 |
| Lineage 4b | 20 | CALD1 | IGFBP3 | 20 | SPARC | CRIM1 |

Pathway analysis revealed significant differences between late-stage lineages from groups (a) and (b). Specifically activated in lineage 1a (group a) were cytokine storm, collagen receptor GP6, and wound healing signaling. IL-4 signaling was also unique to lineage 1, whereas IL-12 activation was observed in both lineage 1a and 3a. Cholesterol signaling was strongly activated in lineage 2a, continuing the expression pattern from middle stage lineage 4. Lineage 2a also upregulated FC gamma-receptor mediated phagocytosis and the NRF2-mediated oxidative stress response. Ferroptosis signaling and CXCR4 activity were increased in lineage 3a, while lineage 4a was characterized by mitochondrial dysfunction, HER-2 signaling, and PAK (p21-activated kinase) signaling. Oxidative phosphorylation activity was high in lineages 1-3a and inhibited in 4a. PKR induction of interferons was activated in lineages 1 and 3 specifically. The second group (b) of late-stage lineages were analyzed separately. We found that lineage 1b specifically activated the unfolded protein response (UPR), the WNT/Ca pathway, mitochondrial dysfunction, and HIPPO and lysosomal signaling. Lineages 1b and 2b upregulated IL-4 signaling and the chaperone mediated autophagy pathway. Interestingly, rheumatoid arthritis signaling was activated specifically in lineage 2b. In lineage 3b, IL-8 and CXCR4 signaling increased, alongside integrin and PI3K/AKT signaling. Finally, lineage 4b shared upregulated EIF2 and erythropoietin signaling with lineage 3b, but uniquely activated NRF2-mediated oxidative stress.

## Discussion

Our results confirm prior reports about the heterogeneity of single SnCs, both within single timepoints and across dynamic senescence development. Far from being a monolithic or static cell state, our data indicates SnCs undergo active transcriptional changes from induction until the last time point at D12. Co-clustering cells from different timepoints showed that upon senescence induction, cells transition to distinct senescent profiles at different speeds. Furthermore, cell lineage trajectories, differentially expressed genes, and differentially regulated pathways showed a wide range of variability in SnC transcriptional activity. Chemokine signaling, mitochondrial and endoplasmic reticulum dysfunction, lysosomal activity, and other altered pathways we identified have previously been associated with SnCs. Our findings at the single-cell RNA level expand on previous work describing the cellular outcomes of doxo induced senescence. Some inducers, such as ionizing radiation (which causes double-stranded DNA breaks), have a singular mechanism by which they induce senescence, which may result in a less variable senescent phenotype. In contrast, our data suggests that doxo induces senescence by multiple mechanisms, including interfering with DNA stability and synthesis (triggering a DNA damage response), disruption of mitochondrial function. In contrast to bulk RNAseq, single cell profiling makes it possible to identify SnC populations originating from different cellular damages. By focusing our analyses on cells from distinct time points, we were able to identify and track specific lineages of SnCs. We describe these lineages in terms of transcriptional state at root, divergence, and end clusters, retracing the transcriptional steps SnCs take as they enter growth arrest. These lineage profiles can be used to predict or manipulate a SnC’s fate, for example to enrich for cells involved in wound healing, or cells characterized by upregulation of specific interleukins. The microfluidics-based single cell processing strategy places some limitations on our data, due to the enlarged and fragile state of SnCs. Some SnCs grow in size, and the largest of these cells may not pass through the channels of the fluidic chips, instead being lysed – releasing the contents of the cells into the ambient medium. Cells also experience shear stresses as they travel through chip, and thus may lyse even before the encapsulation step. As a result, a caveat of our data is that the identified clusters and reconstructed lineages are not fully representative of the doxo-induced SnC pool. These enlarged and/or fragile cells that are not profiled form an understudied population that needs to be studied using different strategies. Solutions to this problem include using nuclei instead of whole cells or using a non-microfluidic approach such as Split-seq. Single nucleus profiling is attractive for several practical reasons such as ease of processing and storage. However the concordance between the transcriptional profile of a whole cell compared to its nucleus varies by cell type. Furthermore, collecting only the nuclear fraction of RNA prevents studying the fully mature population of mRNAs in the cytosol. The Split-seq approach to single cell profiling eschews microfluidics altogether. Instead of droplet encapsulation, fixed and permeabilized cells are used as the vessels for the reverse transcription of mRNA into cDNA. This plate-based system circumvents cell size limitations, and allows for heterogeneous cells to be processed simultaneously. Combined with gentle pipetting, the lysis of fragile cells can also be reduced this way, which may provide a more complete picture of the transcriptomics of senescence.

## Methods

### IMR90 culture and senescence induction

All experiments were performed in incubators at 37 °C, 3% O2 and 5% CO2. Senescence was induced by treatment with doxo (250 nM for 24 h) (Demaria et al., 2017). Proliferating cells were harvested from non-synchronized normally cycling cells in 10% FBS. Quiescence was induced by serum starvation (0.2% FBS) for 3 days, followed by cell collection. Treated cells were harvested at post-induction timepoints including 12 hours post-doxo, 36 hours, and days 2, 4, 6, 8, 10, and 12.

### 10x Single cell library preparation and sequencing

Harvested cells were prepared as input for 10x Chromium Next GEM Single Cell 3’ Reagent v3.1 kits. Sample processing and library preparation was performed according to the 10X supplied protocol. To reduce batch effects, only cell encapsulation and cDNA conversion for each sample was performed on the day of cell harvest. Further processing and library construction were then performed simultaneously for each sample. QC for cDNA and libraries was performed using the Qubit fluorometer and Agilent Tapestation, according to the 10X protocol. Sequencing was performed on an S4 flowcell type Illumina NovaSeq 6000 system.

### Single cell data analysis

Raw sequencing data was processed with Cellranger count (version 3.0.2). Cellranger output was then processed in Seurat (version 4.2). Doublets were identified with the DoubletFinder R package (version 2.0). Cells were excluded from the analysis unless they met the following criteria: percent of reads mapping to MALAT1 < 20, percent reads NEAT1 < 20, nFeature_RNA > 500, nCount_RNA > 2000, percent mitochondrial reads < 25, percent ribosomal reads < 25. SCTransform was used to perform normalization, variance stabilization, and feature selection. Percent reads from mitochondrial, ribosomal, MALAT1, and NEAT1 expression were regressed in a second linear regression. Proliferating cells were identified by expression of MKI67, CCNA2, TOP2A, MCM2, and PCNA. Further details and R scripts used for sample integration, cluster identification, DEG analyses, and trajectory fitting and visualization using Slingshot and tradeSeq, are available in supplemental materials (XXX). IPA core analysis was used to identify differentially regulated pathways and regulatory element activity.

## Notes

### Competing Interest Statement

The authors have declared no competing interest.

## References

1. Bodnar, A.G., et al., Extension of life-span by introduction of telomerase into normal human cells. Science, 1998. 279(5349): p. 349–52.

2. Campisi, J. and F. d’Adda di Fagagna, Cellular senescence: when bad things happen to good cells. Nat Rev Mol Cell Biol, 2007. 8(9): p. 729–40.

3. Gorgoulis, V., et al., Cellular Senescence: Defining a Path Forward. Cell, 2019. 179(4): p. 813–827.

4. Serrano, M., et al., Oncogenic ras provokes premature cell senescence associated with accumulation of p53 and p16INK4a. Cell, 1997. 88(5): p. 593–602.

5. Aguayo-Mazzucato, C., et al., Acceleration of beta Cell Aging Determines Diabetes and Senolysis Improves Disease Outcomes. Cell Metab, 2019. 30(1): p. 129–142 e4.

6. Baker, D.J., et al., Naturally occurring p16(Ink4a)-positive cells shorten healthy lifespan. Nature, 2016. 530(7589): p. 184–9.

7. Bussian, T.J., et al., Clearance of senescent glial cells prevents tau-dependent pathology and cognitive decline. Nature, 2018. 562(7728): p. 578–582.

8. Campisi, J., Cellular senescence as a tumor-suppressor mechanism. Trends Cell Biol, 2001. 11(11): p. S27–31.

9. Childs, B.G., et al., Senescent intimal foam cells are deleterious at all stages of atherosclerosis. Science, 2016. 354(6311): p. 472–477.

10. Chinta, S.J., et al., Cellular Senescence Is Induced by the Environmental Neurotoxin Paraquat and Contributes to Neuropathology Linked to Parkinson’s Disease. Cell Rep, 2018. 22(4): p. 930–940.

11. Demaria, M., et al., Cellular Senescence Promotes Adverse Effects of Chemotherapy and Cancer Relapse. Cancer Discov, 2017. 7(2): p. 165–176.

12. Demaria, M., et al., An essential role for senescent cells in optimal wound healing through secretion of PDGF-AA. Dev Cell, 2014. 31(6): p. 722–33.

13. Farr, J.N., et al., Targeting cellular senescence prevents age-related bone loss in mice. Nat Med, 2017. 23(9): p. 1072–1079.

14. Jeon, O.H., et al., Local clearance of senescent cells attenuates the development of post-traumatic osteoarthritis and creates a pro-regenerative environment. Nat Med, 2017. 23(6): p. 775–781.

15. Menon, R., et al., Placental membrane aging and HMGB1 signaling associated with human parturition. Aging (Albany NY), 2016. 8(2): p. 216–30.

16. Munoz-Espin, D., et al., Programmed cell senescence during mammalian embryonic development. Cell, 2013. 155(5): p. 1104–18.

17. Ohtani, N., D.J. Mann, and E. Hara, Cellular senescence: its role in tumor suppression and aging. Cancer Sci, 2009. 100(5): p. 792–7.

18. Storer, M., et al., Senescence is a developmental mechanism that contributes to embryonic growth and patterning. Cell, 2013. 155(5): p. 1119–30.

19. Kumari, R. and P. Jat, Mechanisms of Cellular Senescence: Cell Cycle Arrest and Senescence Associated Secretory Phenotype. Front Cell Dev Biol, 2021. 9: p. 645593.

20. Hernandez-Segura, A., et al., Unmasking Transcriptional Heterogeneity in Senescent Cells. Curr Biol, 2017. 27(17): p. 2652–2660 e4.

21. van Deursen, J.M., The role of senescent cells in ageing. Nature, 2014. 509(7501): p. 439–46.

22. Acosta, J.C., et al., Chemokine signaling via the CXCR2 receptor reinforces senescence. Cell, 2008. 133(6): p. 1006–18.

23. Coppe, J.P., et al., Senescence-associated secretory phenotypes reveal cell-nonautonomous functions of oncogenic RAS and the p53 tumor suppressor. PLoS Biol, 2008. 6(12): p. 2853–68.

24. Kuilman, T., et al., Oncogene-induced senescence relayed by an interleukin-dependent inflammatory network. Cell, 2008. 133(6): p. 1019–31.

25. Kuilman, T. and D.S. Peeper, Senescence-messaging secretome: SMS-ing cellular stress. Nat Rev Cancer, 2009. 9(2): p. 81–94.

26. Nelson, G., et al., A senescent cell bystander effect: senescence-induced senescence. Aging Cell, 2012. 11(2): p. 345–9.

27. Acosta, J.C., et al., A complex secretory program orchestrated by the inflammasome controls paracrine senescence. Nat Cell Biol, 2013. 15(8): p. 978–90.

28. The gene expression program of prostate fibroblast senescence modulates neoplastic epithelial cell proliferation through paracrine mechanisms.

29. Rodier, F., et al., Persistent DNA damage signalling triggers senescence-associated inflammatory cytokine secretion. Nat Cell Biol, 2009. 11(8): p. 973–9.

30. Novakova, Z., et al., Cytokine expression and signaling in drug-induced cellular senescence. Oncogene, 2010. 29(2): p. 273–84.

31. Aging, cellular senescence, and cancer.

32. Niedernhofer, L.J. and P.D. Robbins, Senotherapeutics for healthy ageing. Nat Rev Drug Discov, 2018. 17(5): p. 377.

33. Chang, J., et al., Clearance of senescent cells by ABT263 rejuvenates aged hematopoietic stem cells in mice. Nat Med, 2016. 22(1): p. 78–83.

34. Hall, B.M., et al., p16(Ink4a) and senescence-associated beta-galactosidase can be induced in macrophages as part of a reversible response to physiological stimuli. Aging (Albany NY), 2017. 9(8): p. 1867–1884.

35. Wiley, C.D., et al., Analysis of individual cells identifies cell-to-cell variability following induction of cellular senescence. Aging Cell, 2017. 16(5): p. 1043–1050.

